# poly(UG)-tailed RNAs in Genome Protection and Epigenetic Inheritance

**DOI:** 10.1101/2019.12.31.891960

**Authors:** Aditi Shukla, Jenny Yan, Daniel J. Pagano, Anne E. Dodson, Yuhan Fei, Josh Gorham, J.G. Seidman, Marvin Wickens, Scott Kennedy

## Abstract

Mobile genetic elements threaten genome integrity in all organisms. MUT-2/RDE-3 is a ribonucleotidyltransferase required for transposon silencing and RNA interference (RNAi) in *C. elegans*. When tethered to RNAs in heterologous expression systems, RDE-3 can add long stretches of alternating non-templated uridine (U) and guanosine (G) ribonucleotides to the 3’ termini of these RNAs (polyUG or pUG tails). Here, we show that, in its natural context in *C. elegans*, RDE-3 adds pUG tails to transposon RNAs, as well as to targets of RNAi. pUG tails with more than 16 perfectly alternating 3’ U and G nucleotides convert otherwise inert RNA fragments into agents of gene silencing. pUG tails promote gene silencing by recruiting RNA-dependent RNA Polymerases (RdRPs), which use pUG-tailed RNAs as templates to synthesize small interfering RNAs (siRNAs). Cycles of pUG RNA-templated siRNA synthesis and siRNA-directed mRNA pUGylation underlie dsRNA-directed transgenerational epigenetic inheritance in the *C. elegans* germline. Our results show that pUG tails convert RNAs into transgenerational memories of past gene silencing events, which, we speculate, allow parents to inoculate progeny against the expression of unwanted or parasitic genetic elements.

## Main text

Transposable elements are mobile parasitic genetic elements present in all genomes. Transposons threaten genome integrity, and can cause disease by disrupting genes or inducing non-allelic recombination. RNA interference (RNAi) is a conserved gene silencing process initiated by double-stranded RNA (dsRNA)^1^. Forward genetic screens to identify factors required for either transposon silencing or RNAi have been conducted in the model metazoan *C. elegans*^2–4^. These screens identified a similar set of genes, indicating that an RNAi-related process silences transposons^2–4^. One gene required for both efficient transposon silencing and RNAi in *C. elegans* is *mut-2/rde-3*, which encodes a protein with homology to ribonucleotidyltransferases (rNTs)^2–5^. rNTs add non-templated ribonucleotides to RNAs and other substrates^6,7^. Recently, *C. elegans* MUT-2/RDE-3 (henceforth, RDE-3) was shown to add perfectly alternating U and G ribonucleotides to the 3’ termini of RNAs (termed polyUG or pUG tails) to which it was tethered in either *S. cerevisiae* or *X. laevis* oocytes^8^. These data suggest that RDE-3 may append non-templated pUG tails to the 3’ termini of RNAs during transposon silencing and/or RNAi in *C. elegans*.

### RNAi directs RDE-3-dependent mRNA pUGylation

We first asked whether pUG tails are added to RNAs targeted by RNAi in *C. elegans*. We exposed animals to dsRNA targeting the germline-expressed gene *oma-1*^9^. Total RNA was extracted and reverse transcribed (RT) using an (AC)_9_ oligo, and nested PCR was used to try to detect *oma-1* RNAs modified with 3’ pUG repeats (Fig. 1a). This approach detected PCR products that were dependent on *oma-1* dsRNA (Fig. 1b), as well as on components of the RNAi machinery including RDE-4, which promotes dsRNA processing into siRNAs^10,11^, the siRNA-binding Argonaute (AGO) protein RDE-1^10,11^, and RDE-8, which is an endonuclease thought to cleave mRNAs exhibiting homology to siRNAs^12^ (Fig. 1c).

**Figure 1.**
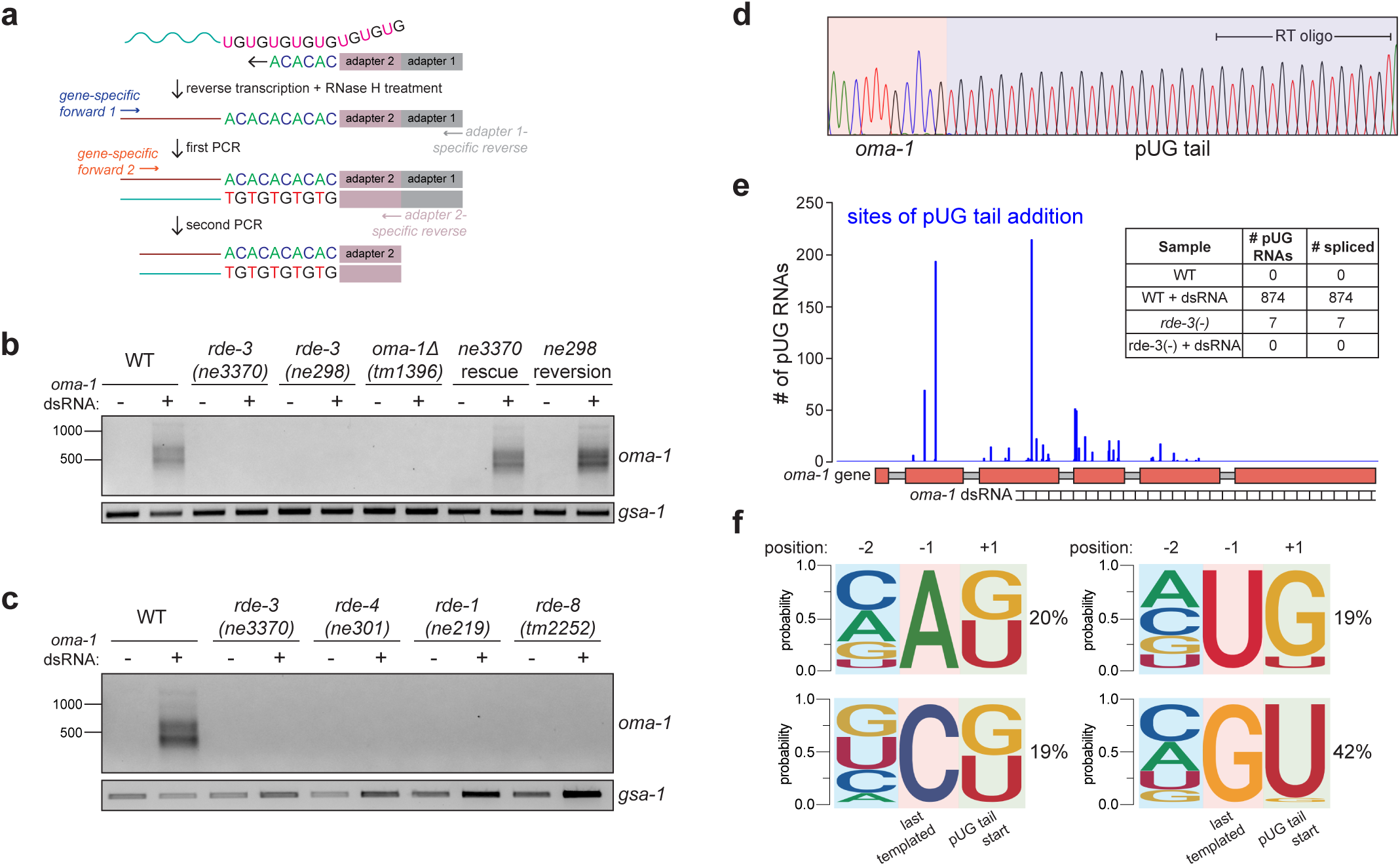
pUG tails are added to mRNA fragments *in vivo*. **a**, PCR-based assay to detect gene-specific pUG RNAs. Total RNA was reverse transcribed (RT) using an (AC)_9_ oligo modified with two PCR adapters and then degraded using RNase H. Two rounds of PCR were performed using gene-specific and adapter-specific primers. Note: the (AC)_9_ RT oligo can complementary base-pair anywhere along the length of a pUG tail. **b**, *oma-1* pUG PCR was performed on total RNA isolated from wild-type animals and two different *rde-3* mutant strains fed *E. coli* expressing empty vector control or *oma-1* dsRNA (RNAi). RDE-3 function was rescued in *ne3370* and *ne298* animals (see Main text and Methods for details). *gsa-1*, which has an 18nt long genomically-encoded pUG repeat in its 3’UTR, is a loading control. **c,** *oma-1* pUG PCR on RNA isolated from animals of the indicated genotypes, +/- *oma-1* dsRNA. **d,** Sanger sequencing chromatogram of an *oma-1* pUG PCR product showing that a pUG tail consists of perfect UG repeats and is longer than the RT oligo. **e,** Illumina MiSeq was performed on *oma-1* pUG PCR products derived from wild-type and *rde-3(-)* animals +/- *oma-1* dsRNA. # of sequenced pUG RNAs (y-axis) mapping to each pUGylation site (x-axis) is shown. Inset: total number of sequenced and spliced *oma-1* pUG RNAs from indicated samples. **f,** % of *oma-1* pUG RNAs (MiSeq reads) having each nucleotide (nt) at the last templated position (−1) is indicated. Logo analysis was used to determine the probability of finding each nt at both the first position of a pUG tail (+1), as well as at the second-to-last templated nt of *oma-1* (−2).

Sanger and Illumina sequencing revealed that most (>89%) pUG PCR products were derived from hybrid RNAs consisting of nearly perfectly alternating (error rate <2%, Table S1) U and G nucleotide (nt) repeats appended to the 3’ termini of sense and spliced *oma-1* mRNA fragments (Fig. 1d, e). Critically, most (64%) pUG tails were longer (range=19-75nt) than the (AC)_9_ oligo used for RT (Fig. 1d, Table S1), indicating that the pUG RNAs we detected were not the result of priming off genomically-encoded UG-rich sequences. Sequencing showed that pUG tails could be appended to any nucleotide of the *oma-1* mRNA; however, if the last templated nucleotide of the *oma-1* mRNA was a G or U, then pUG tails tended to initiate with U (96%) or G (88%), respectively (Fig. 1f). RDE-3 was required for addition of pUG repeats to mRNA fragments (termed pUGylation): *rde-3* mutants, including *rde-3(ne3370)* animals, which harbor a deletion that removes residues required for catalysis within the rNT domain of RDE-3 (henceforth *rde-3(-)*)^8^, failed to produce *oma-1* pUG RNAs in response to *oma-1* dsRNA (Fig. 1b). pUGylation defects in *rde-3* mutants were rescued by introducing a wild-type copy of *rde-3* into *rde-3(-)* animals or by CRISPR/Cas9-mediated reversion of a missense allele (*ne298)* of *rde-3* to wild-type (Fig. 1b). RNA pUGylation was a general response to RNAi: animals exposed to dsRNA targeting a germline-expressed *gfp::h2b* transgene or the hypodermally-expressed *dpy-11* mRNA^13^ produced RDE-3-dependent *gfp* and *dpy-11* pUG RNAs, respectively (Extended Data Fig. 1). Furthermore, pUGylation was sequence-specific, since *dpy-11* dsRNA did not induce *oma-1* pUG RNA biogenesis and vice versa (Extended Data Fig. 1). Together, these data indicate that RDE-3 adds pUG tails to mRNAs targeted for silencing by RNAi.

### pUG tails turn inert mRNA fragments into agents of gene silencing

Given that RNAi is a gene-silencing phenomenon^1^, pUG tails could either mark mRNA fragments for degradation or convert mRNA fragments into active mediators of gene silencing. To differentiate these possibilities, we asked whether *in vitro* transcribed pUG RNAs possessed gene silencing activity. Indeed, injection of a *gfp* pUG RNA, consisting of 18 3’terminal pUG repeats appended to the first 369nts of the *gfp* mRNA, into animals expressing a germline-expressed *gfp::h2b* transgene was sufficient to silence *gfp::h2b* expression (Fig. 2a). The same *gfp* mRNA fragment without a 3’ tail or with 18 3’-terminal pGC, pAC, or pAU repeats lacked gene silencing activity (Fig. 2a). Note: to control for potential dsRNA contamination in our *in vitro* transcription reactions, all RNAs were injected into *rde-1(ne219)* mutant animals, as *rde-1* mutants cannot respond to dsRNA (Fig. 2a, b)^4^. The ability of a pUG tail to turn an mRNA fragment into an agent of gene silencing was both general and sequence-specific. *oma-1(zu405ts)* animals lay arrested embryos unless *oma*-*1(zu405ts)* is silenced^14^. An *in vitro* transcribed 541nt long *oma-1* mRNA fragment modified with 18 3’ pUG—but not 18 pGC, pAC, or pAU—repeats was capable of silencing *oma-1(zu405ts)* (Fig. 2b). Additionally, an *oma-1* pUG RNA injection did not silence *gfp::h2b* and *vice versa* (Extended Data Fig. 2). We conclude that pUG tails convert otherwise inert mRNA fragments into agents of gene silencing.

**Figure 2.**
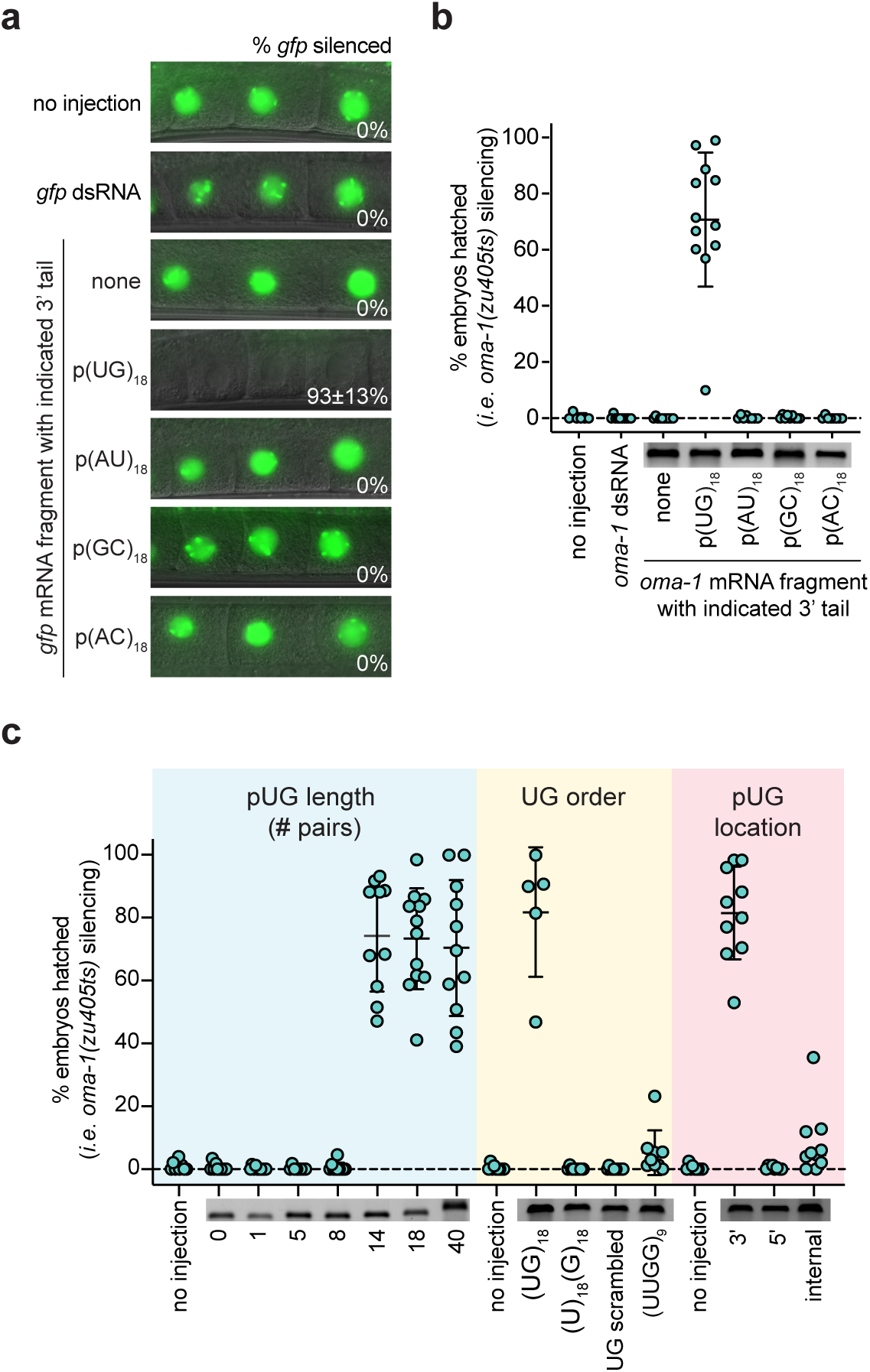
pUG tails convert otherwise inert RNAs into agents of gene silencing. **a**, Fluorescent micrographs showing -1 to -3 oocytes of adult *rde-1(ne219); gfp::h2b* animals injected in the germline with RNAs consisting of the indicated 3’ terminal repeats appended to the first 369nt of *gfp* mRNA. % of progeny with *gfp* silenced was counted. **b,** *oma-1(zu405ts)* animals lay arrested embryos at 20°C unless *oma-1(zu405ts)* is silenced^14^. Adult *rde-1(ne219); oma-1(zu405ts)* animals were injected with RNAs consisting of the indicated 3’ terminal repeats appended to the first 541nt of *oma-1* mRNA. **c,** Adult *rde-1(ne219); oma-1(zu405ts)* animals were injected with the same *oma-1* mRNA fragment as in **b** with varying 3’ pUG tail length, different UG repeat sequences or with the pUG sequence appended to the 3’ end, 5’ end or in the middle of the *oma-1* mRNA. **b-c,** 5 progeny per injected animal were pooled and the % hatched embryos (# of hatched embryos/total embryos laid) was counted. Insets show injected RNAs run on a 2% agarose gel to assess RNA integrity. n=5-15 injected animals. **a-c,** Repeats were 36nt in length unless otherwise indicated. Error bars are standard deviations (s.d.) of the mean.

We used the pUG RNA injection assay to define the features of pUG RNAs required for biological activity. We injected animals with *oma-1* pUG RNAs harboring varying numbers of 3’ UG repeats and found that 14, 18, or 40—but not 1, 5, or 8—UG repeats were capable of triggering *oma-1* gene silencing when appended to the same *oma-1* mRNA fragment (Fig. 2c). We also found that while pUG tails with perfectly alternating U and G nucleotide repeats conferred silencing activity to an mRNA fragment, 3’ tails with scrambled UG sequence or other combinations of Us and Gs did not (Fig. 2c). Moreover, while an *oma-1* mRNA fragment with a 3’ pUG tail triggered *oma-1(zu405ts)* silencing, *oma-1* mRNA fragments with 5’ or internal UG repeats did not (Fig. 2c). Finally, the *oma-1* segment of an *oma-1* pUG RNA had to possess the sense coding sequence and be >50nts in length for pUG RNA functionality (Extended Data Fig. 3). Together, these data show that a pUG RNA must consist of >8 3’ UG repeats appended to >50nt of sense RNA in order to trigger gene silencing.

### RDE-3 pUGylates germline-expressed RNAs

We next asked whether endogenous mRNAs are pUGylated in *C. elegans*. In the absence of RDE-3, Tc1 transposase RNA is upregulated and Tc1 mobilizes^2^, suggesting that the Tc1 RNA might be pUGylated. We tested this idea using a Tc1-specific pUG PCR assay (Fig. 1a) and observed RDE-3–dependent pUG tails appended to Tc1 RNA fragments, which were between 41-195nt in length (Fig. 3a and Table S1). In addition, Tc1 mobilization caused by *rde-3* mutation was suppressed by injection of a Tc1 pUG RNA (Fig. 3b). We conclude that RDE-3–based pUGylation silences the Tc1 transposon in *C. elegans*.

**Figure 3.**
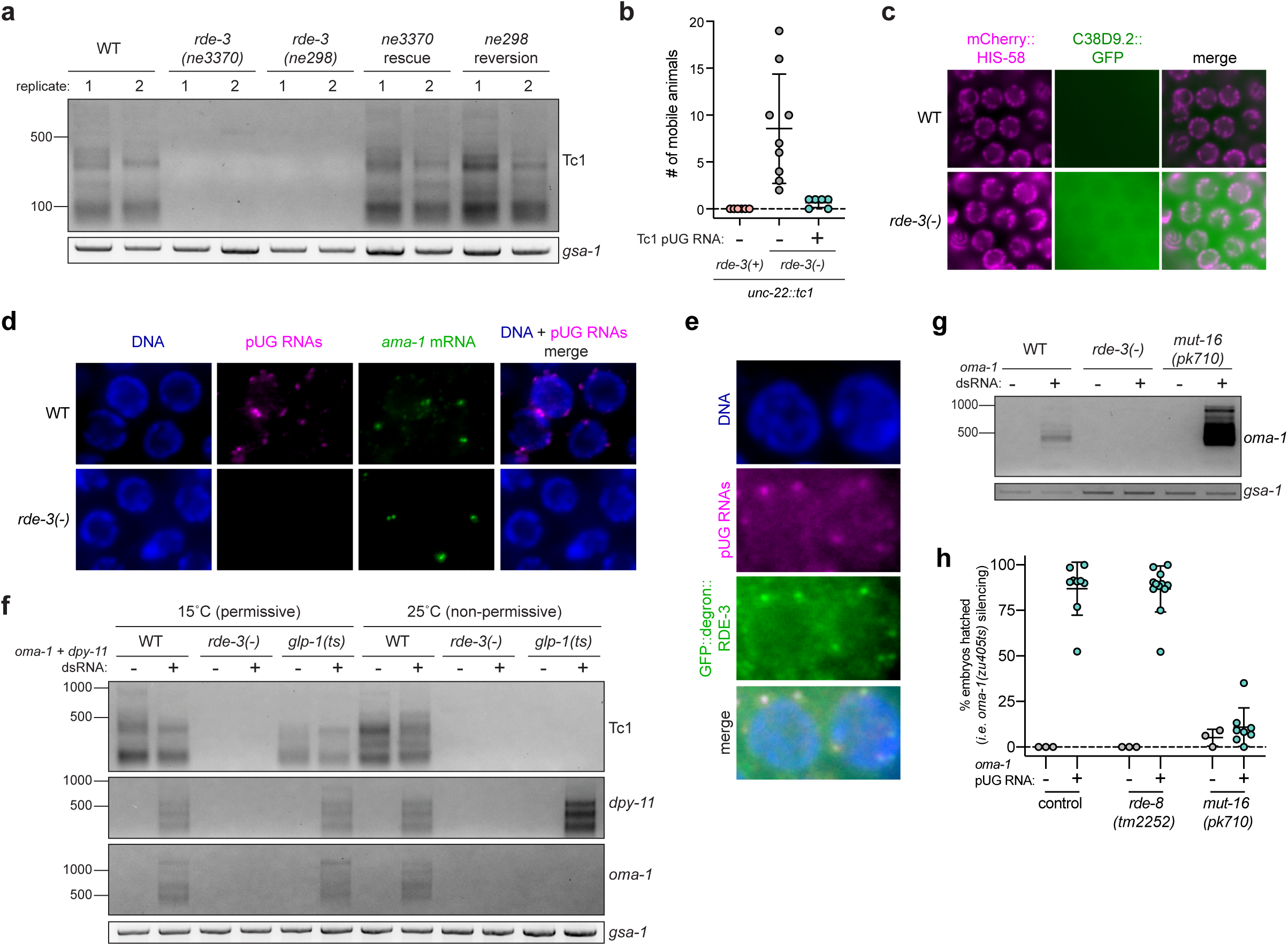
Endogenous RNAs are pUGylated and localize to germline *Mutator* foci. **a**, Total RNA isolated from adult wild-type or *rde-3* mutant animals was subjected to Tc1 pUG PCR analysis (Fig. 1a). Rescue strategies are described in the Main text and Methods. **b,** A 36nt pUG tail was appended to a 338nt Tc1 RNA fragment and this Tc1 pUG RNA was injected into germlines of *rde-3(-); unc-22::tc1* animals with a co-injection marker. 25 co-injection marker expressing progeny were pooled per injected animal. Each data point represents the # of mobile progeny (indicating Tc1 mobilized from *unc-22*) per pool. Error bars represent s.d. **c-e,** Fluorescent micrographs of adult pachytene germ cell nuclei. **c,** Wild-type or *rde-3(-)* animals expressing a marker of chromatin (mCherry:HIS-58, magenta) and C38D9.2::GFP (green), which is expressed diffusely in the germline syncytium, wherein germ cell nuclei share a common cytoplasm. **d,** RNA FISH to detect pUG RNAs (pUG RNA FISH) was performed on germlines dissected from wild-type or *rde-3(-)* animals using an 18nt long poly(AC) oligo conjugated to Alexa 647 (magenta). RNA FISH to detect *ama-1* mRNA (green) was performed simultaneously as a positive control. DNA was stained with DAPI (blue). **e,** pUG RNA FISH (magenta) and immunofluorescence to detect a GFP- and degron-tagged RDE-3 (green). DNA was stained with DAPI (blue). **f,** Tc1, *dpy-11*, and *oma-1* pUG PCR was performed on total RNA isolated from *glp-1(q224* or *ts)* animals grown at 15°C (permissive temperature, germ cells present) or 25°C (non-permissive temperature, <99% of germ cells) fed empty vector control or *oma-1* and *dpy-11* dsRNA simultaneously. **g,** *oma-1* pUG PCR was performed on total RNA extracted from wild-type, *rde-3(-)*, and *mut-16(pk710)* animals +/- *oma-1* dsRNA. Note: pUG RNAs appear longer in *mut-16* mutants (see Extended Data Fig. 9 legend). **h,** Control, *rde-8(tm2252)* or *mut-16(pk710)* animals (all *rde-1(ne219); oma-1(zu405ts)* background) were injected with *oma-1* pUG RNAs and % embryos hatched was scored. Error bars +/- s.d. n=8-12 injected animals.

To identify additional targets of pUGylation, we conducted mRNA-seq on wild-type and *rde-3(-)* animals and identified 346 RNAs that were upregulated in *rde-3(-)* animals (Table S2, adjusted p value <0.05 and log2 fold change >1.5). We observed increased levels of Tc1 RNA, as well as six other DNA transposons (Tc1A, TC4, TC5, MIRAGE1, CEMUDR1, Chapaev-2), several LTR retrotransposons (CER3, CER9, CER13), and 294 predicted protein-coding RNAs (Table S2). Directed pUG RT-PCR analyses confirmed that Tc4v, Tc5, CER3, and four of five genes tested from amongst our list of top 25 most RDE-3–regulated mRNAs, were pUGylated in an RDE-3 dependent manner (Extended Data Fig. 4). pUG tails were not detected on RNAs whose expression is unchanged in *rde-3* mutants, such as *oma-1, gfp* or *dpy-11* (Extended Data Fig. 1) and two additional genes selected at random (Extended Data Fig. 4). We used CRISPR/Cas9 to introduce a *gfp* tag into one RDE-3–regulated and pUGylated locus, *c38d9.2*. We observed diffuse C38D9.2::GFP expression in the germline syncytium of *rde-3(-)*, but not *rde*-*3(+)*, animals, confirming that *c38d9.2* is regulated by RDE-3 (and thus, pUGylation) and showing that this regulation occurs in the germline (Fig. 3c). We conclude that RDE-3 adds pUG tails to endogenous RNAs in *C. elegans*, which include, but are not limited to, transposon RNAs.

### pUG RNAs and germ granules

Germ granules are liquid-like condensates that form near the outer nuclear membrane in most animal germ cells^15^. Germ granules are thought to promote germ cell totipotency by concentrating germline determinants, which include maternal RNAs and proteins, into developing germline blastomeres^15^. *C. elegans* RDE-3 localizes to perinuclear germ granules termed *Mutator* foci^16^. RNA fluorescence *in situ* hybridization (RNA FISH) using a fluorescently labeled p(AC)_9_ probe to detect pUG RNAs (pUG FISH) showed that pUG RNAs localized to perinuclear puncta in germ cells of *rde-3(+)*, but not *rde-3(-),* animals (Fig. 3d). pUG FISH coupled with immunofluorescence (IF) to detect a GFP- and degron-tagged RDE-3 showed that pUG RNA foci co-localized with RDE-3 and, therefore, *Mutator* foci (Fig. 3e). This data suggests that pUG RNAs are produced, function, and/or stored in *Mutator* foci in the *C. elegans* germline. Indeed, *glp-1(q224)* animals, which lack ≅99% of their germ cells when grown at 25°C (hereafter, *glp-1(ts))*, failed to produce detectable Tc1 pUG RNAs (or *oma-1* dsRNA-induced *oma-1* pUG RNAs) when grown at 25°C, confirming that pUG RNAs are produced or stored in germ cells (Fig. 3f)^17^. Incidentally, a related analysis with *glp-1(ts)* animals treated with dsRNA targeting the hypodermally-expressed *dpy-11* gene^13^ showed that pUG RNAs can also be produced in somatic cells (Fig. 3f). Hereafter, however, we focus on exploring the biogenesis and function of pUG RNAs in the germline.

To explore further how germline pUG RNAs and *Mutator* foci might relate, we asked if the glutamine/asparagine(Q/N) motif-rich protein MUT-16, which is required for *Mutator* foci assembly in germ cells^16^, was needed for pUG RNA biogenesis or function. *mut-16(pk710)* animals, which harbor a nonsense mutation in *mut-16,* produced *oma-1* pUG RNAs in response to *oma-1* dsRNA (Fig. 3g), but failed to silence *oma-1* after an *oma-1* pUG RNA injection (Fig. 3h). These data suggest that *Mutator foci* are required for pUG RNA-based gene silencing, downstream of pUG RNA biogenesis.

### pUG tails convert RNAs into templates for RNA-dependent RNA Polymerases (RdRPs)

RdRPs amplify RNAi-triggered gene silencing signals in *C. elegans*^18^. Current models posit that RdRPs: 1) are recruited to mRNAs by siRNAs generated from dsRNA, and 2) use these mRNAs as templates to synthesize additional siRNAs, termed secondary (2°) siRNAs, which carry out gene silencing^19–21^. RRF-1, one of the four *C. elegans* RdRPs, localizes to *Mutator* foci^16^. We wondered whether pUG tails might promote gene silencing by recruiting RdRPs, such as RRF-1, to pUG RNAs, which could then act as templates for siRNA synthesis. To first ask if the pUG tail of a pUG RNA is sufficient to recruit RRF-1, we conjugated 5’ biotinylated RNAs consisting of 5, 8, 14, or 18 UG repeats; 18 GC repeats; or 36 scrambled UGs to streptavidin beads and incubated these beads with extracts obtained from animals expressing HA::TagRFP::RRF-1. α-HA immunoblotting showed that HA::TagRFP::RRF-1 pelleted with (UG)_18_, but not (GC)_18_ or UG scrambled RNAs (Fig. 4a). Additionally, HA::TagRFP::RRF-1 pelleted strongly with (UG)_14_ and (UG)_18_ RNAs, weakly with a (UG)_8_ RNA, and not with a (UG)_5_ RNA (Fig. 4b). These data show that the RdRP RRF-1 interacts physically with UG repeat RNAs and that the sequence determinants of this interaction largely mirror those required for pUG tail-mediated gene silencing *in vivo* (Fig. 2c).

**Figure 4.**
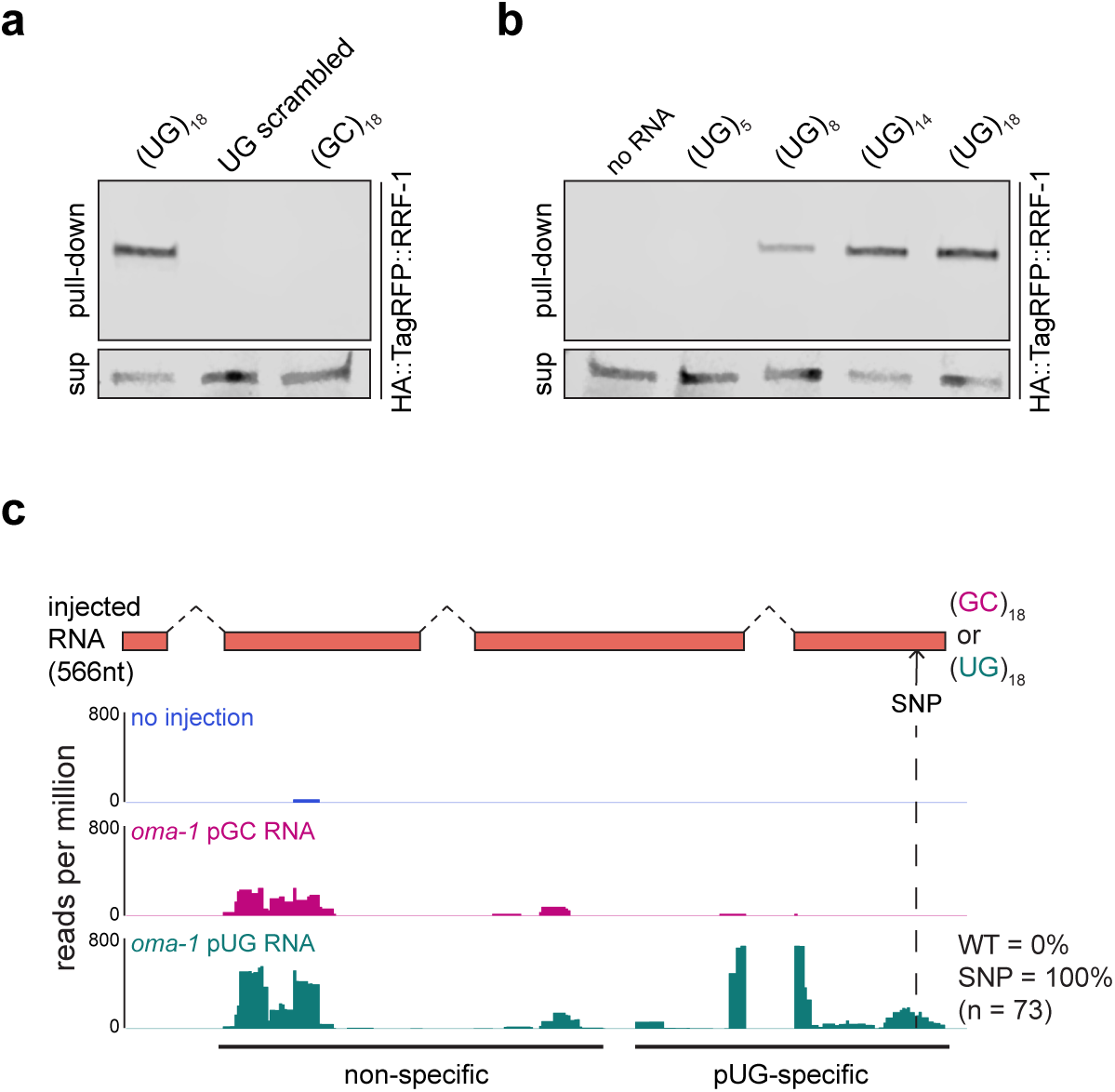
pUG RNAs are templates for RdRPs. **a-b**, The indicated 5’ biotinylated RNA oligos were conjugated to streptavidin beads. Beads were then incubated with extracts generated from animals expressing an HA- and tagRFP-tagged RRF-1 and pelleted to separate beads (pull-down) from supernatant (sup). Both pull-down and sup fractions were subjected to α-HA immunoblotting. **c,** *rde-1(ne219); oma-1(zu405ts)* animals were injected with a SNP-containing (dotted line) *oma-1 (oma-1(SNP))* pUG or pGC RNA. Injected animals were collected 1-4 hours after injection and small RNAs (20-30nts) were sequenced. In *C. elegans,* RdRP-derived small RNAs are antisense, 22nt in length, and initiate with guanosine (termed secondary siRNAs or 22G siRNAs)^22^. 22G siRNAs mapping antisense to *oma-1* are shown. Injection of an *oma-1(SNP)* pUG (but not pGC) RNA triggered 22G siRNA production near the site of the pUG tail (pUG-specific). 100% of these pUG-specific 22G siRNAs contained the engineered SNP. Both *oma-1(SNP)* pUG and pGC RNA injections triggered small RNA production 5’ of the pUG tail (non-specific). The origin of non-specific siRNAs is not known. For a list of all small RNAs sequenced, see Table S3.

To determine whether pUG RNAs act as templates for RdRPs *in vivo*, we sequenced small (20-30nts) RNAs from animals injected with either an *oma-1* pGC or pUG RNA engineered to contain a single-nucleotide polymorphism (SNP) not present in the genomic copy of *oma-1*. This SNP enabled differentiation of siRNAs templated from genomically-encoded *oma-1* mRNAs versus those templated from injected *oma-1(SNP)* pGC or pUG RNAs. In *C. elegans,* RdRP-derived (2°) siRNAs are also known as 22G siRNAs as they are typically antisense, 22nt in length and begin with a guanosine^22^. Small RNA sequencing showed that injection of the *oma-1(SNP)* pUG RNA, but not the *oma-1(SNP)* pGC RNA, triggered the synthesis of *oma-1* 22G siRNAs mapping near (≅100bp upstream) the site where the pUG tail was appended (Fig. 4c, Extended Data Fig. 5 and Table S3)^23^. For unknown reasons, both *oma-1(SNP)* pUG and pGC RNAs triggered non-pUG-specific siRNA synthesis ≅0.4kb upstream of where the tails were appended. Importantly, most (92-100%) pUG-specific 22G siRNAs encoded the complement of the engineered SNP, indicating that these siRNAs were templated from the injected *oma-1(SNP)* pUG RNA (Fig. 4c and Extended Data Fig. 5). We conclude that one function of a pUG tail is to convert RNAs into templates for RdRPs.

### pUG tails convert RNAs into vectors of transgenerational epigenetic inheritance (TEI)

RNAi-triggered gene silencing can be inherited for multiple generations in *C. elegans,* making RNAi inheritance a robust and dramatic example of TEI^24–29^. Interestingly, a one-time exposure of animals to *oma-1* dsRNA not only initiated the production of *oma-1* pUG RNAs, but also caused *oma-1* pUG RNAs to be expressed for four additional generations (Fig. 5a), concomitant with *oma-1* gene silencing (Extended Data Fig. 6). Thus, pUG RNA expression correlates with gene silencing during TEI, consistent with the idea that pUG RNAs may contribute in some way to TEI. To test this idea, we injected animals with *gfp* or *oma-1* pUG RNAs and monitored *gfp* or *oma-1* silencing over generations. *gfp* or *oma-1* pUG RNAs were sufficient to silence *gfp* or *oma-1*, respectively, for multiple generations (Fig. 5b and Extended Data Fig. 7). We conclude that pUG RNAs can induce TEI.

**Figure 5.**
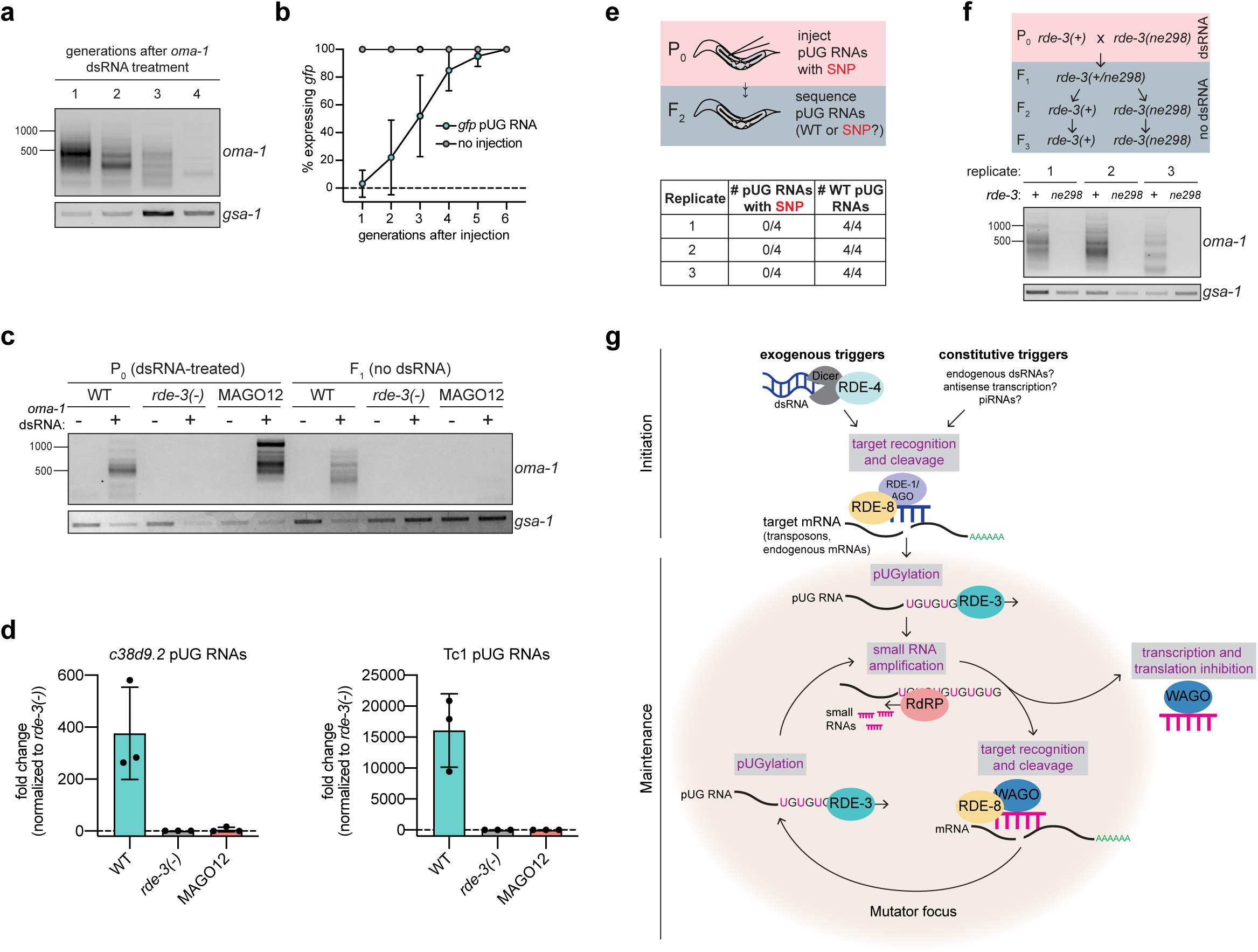
pUG RNAs and siRNAs cooperate to drive heritable gene silencing. **a**, *oma-1* pUG PCR was performed on RNA isolated from four generations of descendants (F_1_-F_4_) derived from *oma-1* dsRNA-treated animals. **b,** *rde-1(ne219); gfp::h2b* animals were injected with a *gfp* pUG RNA and *gfp* expression was monitored for six generations. **c,** MAGO12 animals, which harbor deletions in all twelve *wago* genes, were treated with *oma-1* dsRNA. *oma-1* pUG PCR was performed on total RNA from dsRNA-treated animals (P_0_) and their progeny (F_1_). Note: pUG RNAs appear longer in MAGO12 animals (see Extended Data Fig. 9 legend). **d,** *c38d9.2* and Tc1 pUG RNA expression levels were quantified in embryos harvested from wild-type, MAGO12, or *rde-3(-)* animals. Shown is the fold change normalized to *rde-3(-).* **e,** *rde-1(ne219); oma-1(zu405ts)* animals were injected with an *oma-1(SNP)* pUG RNA. pUG RNAs were Sanger sequenced from F_2_ progeny to determine the presence or absence of the SNP. **f,** Wild-type and *rde-3(ne298)* animals subjected to *oma-1* RNAi were crossed and F_2_ progeny were genotyped (not shown). RNA isolated from populations of *rde-3(+)* or *rde-3(ne298)* F_3_ animals (3 biological replicates) was subjected to *oma-1* pUG PCR. **g,** Model. Two major phases of the pUG RNA pathway, initiation and maintenance, are shown. *Initiation*: exogenous and constitutive (i.e. genomically-encoded such as dsRNA, piRNAs) triggers direct RDE-3 to pUGylate RNAs previously fragmented by RNAi, and possibly other, systems. *Maintenance*: pUG RNA are templates for RdRPs to make 2° siRNAs. Argonaute proteins (termed WAGOs) bind these 2° siRNAs and: 1) target homologous RNAs for transcriptional and translational silencing (previous work^25,30,38,39^), and 2) direct the cleavage and *de novo* RDE-3-mediated pUGylation of additional mRNAs (this work). In this way, cycles of pUG RNA-based siRNA production and siRNA-directed mRNA pUGylation form a silencing loop, which is maintained over time and across generations to mediate stable gene silencing. pUG/siRNA cycling likely occurs in germline perinuclear condensates called *Mutator* foci.

How might pUG RNAs drive TEI? We speculated that if pUG RNA-templated siRNAs (Fig. 4c and Extended Data Fig. 5) could direct *de novo* mRNA pUGylation, then generationally repeated cycles of pUG RNA-templated siRNA synthesis and siRNA-directed pUG RNA biogenesis could be maintained in the absence of initiating dsRNA triggers and, thus, drive TEI. Three lines of evidence support this “pUG/siRNA cycling” model for RNAi-directed TEI. First, *C. elegans* 2° siRNAs can engage at least twelve AGO proteins (termed WAGOs) to mediate gene silencing^30^. MAGO12 animals, which lack all twelve of these WAGOs, produced *oma-1* pUG RNAs after *oma-1* RNAi (Fig. 5c). Progeny of RNAi-treated MAGO12 animals, however, did not inherit the ability to produce *oma-1* pUG RNAs (Fig. 5c). Thus, the 2° siRNA system is needed to maintain pUG RNA expression specifically during the inheriting generations of TEI, which supports a pUG/siRNA cycling model for TEI. Interestingly, pUG RNAs derived from endogenous pUGylation targets *c38d9.2* and Tc1 were also dependent upon the WAGOs (Fig. 5d), suggesting that the endogenous targets of RDE-3 also undergo heritable silencing via pUG/siRNA cycling in the germline.

Second, when we injected animals with an *oma-1(SNP)* pUG RNA, pUG RNAs were detectable in subsequent generations (Extended Data Fig. 8); however, these pUG RNAs did not contain the engineered SNP (Fig. 5e). Similarly, <1% of siRNAs sequenced from progeny of *oma-1(SNP)* pUG RNA injected animals possessed the SNP complement (Extended Data Fig. 8). Combined, these data show that most pUG RNAs and 2° siRNAs expressed during the inheriting generations of RNAi-directed TEI result from *de novo* pUGylation events, supporting the idea that repeated pUG/siRNA cycling mediates TEI.

Third, we conducted a genetic analysis that showed that *de novo* pUGylation events in progeny were required for TEI. We crossed *oma-1* RNAi-treated animals with *rde-3(ne298)* males, isolated *rde-3(+)* and *rde-3(ne298)* F_2_ progeny, and then assayed the F_3_ generation of this cross for *oma-1* pUG RNA expression and *oma-1* gene silencing (Fig. 5f). *rde-3(ne298)* animals failed to express *oma-1* pUG RNAs (Fig. 5f) or silence the *oma-1* locus (Extended Data Fig. 8) during the inheriting generations of TEI, supporting the idea that pUG RNA biogenesis and, therefore, pUG/siRNA cycling in progeny is necessary for TEI maintenance. We conclude that pUG tails convert otherwise inert RNA fragments into drivers of a transgenerational RNA-based memory system, which is likely propagated across generations via iterative cycles of sense pUG RNA and antisense siRNA biogenesis.

## Discussion

Here, we show that RDE-3 adds 3’ UG repeats to germline and soma-expressed RNAs in *C. elegans*, revealing a previously unknown form of RNA modification *in vivo*. RdRPs are recruited to pUG tails and use pUG RNAs, heretofore unrecognized RNA intermediates in the RNAi pathway, as templates for siRNA synthesis (Fig. 5g). Functional pUG tails consist of more than eight pairs of perfect or near-perfect 3’ UG repeats. The precise sequence requirements for pUG tail function hint that pUG tails may form a structure which helps to recruit, and possibly prime, RdRPs. pUG tails/structures might also act as binding sites for other proteins, such as the *C. elegans* ortholog of the mammalian pUG binding protein TDP-43, to regulate the localization, stability, or function of pUG-tailed RNAs.

pUG RNAs act as informational vectors for TEI when they engage in feed-forward amplification cycles with RdRP-generated 2° siRNAs (Fig. 5g). These pUG/siRNA cycles, we speculate, allow *C. elegans* to remember past gene silencing events and inoculate progeny against expressing unwanted and/or dangerous genetic elements. pUG/siRNA cycling likely occurs in *Mutator* foci, germline condensates whose assembly we find is required for pUG/siRNA pathway function (Fig. 5g). Experimental RNAi-initiated pUG/siRNA cycles perdure for several generations, but are not permanent, suggesting that *C. elegans* possess systems to prevent pUG/siRNA cycles from propagating in perpetuity. Interestingly, we find that RNAi-initiated pUG RNAs shorten progressively during TEI (Fig. 5a), suggesting that pUG RNA shortening, which may be an inevitable consequence of RdRP-based 2° siRNA synthesis (Extended Data Fig. 9), could function as one such brake on TEI. In contrast, the natural targets of pUGylation, such as transposons, are constitutively silenced by the pUG/siRNA system, suggesting that genetic systems, such as genomically-encoded PIWI-interacting (pi)RNAs or endogenous dsRNAs, are likely to reinforce and refocus epigenetic pUG/siRNA silencing at these loci each generation (Fig. 5g).

The logic of the *C. elegans* sense-antisense pUG/siRNA TEI system resembles that of fly and mammalian piRNA “ping-pong” systems in which iterative base-pairing between genomically-encoded sense-antisense transposon RNAs, as well as piRNAs derived from these RNAs, mediates stable transposon silencing^31^. We speculate that related sense-antisense RNA systems could contribute to other biological processes during which long-term memories of past expression states are needed, such as development or inheritance of environmentally-triggered acquired traits^32–37^. Finally, our data show that long non-templated and non-homopolymeric tracts of ribonucleotides can be appended to, and confer novel functions to, RNAs in *C. elegans*. It will be of obvious interest to ask whether pUG-tailed RNAs, or RNAs bearing other unexpected tails, are restricted to *C. elegans* or are, instead, emissaries of a new class of eukaryotic RNAs.

## Extended Data Figure Legends

**Extended Data Fig. 1.**
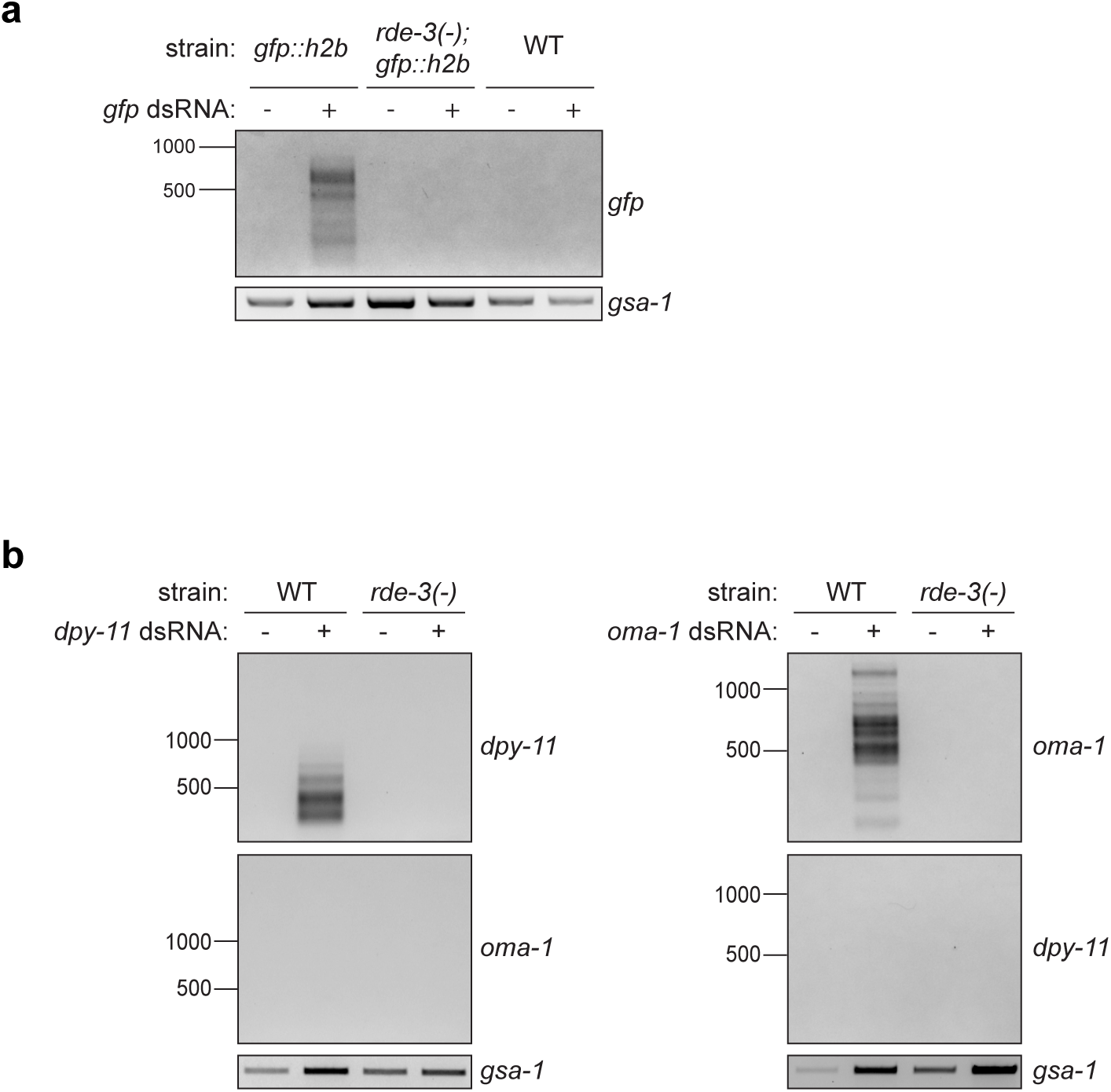
RNAi-triggered pUGylation is general and sequence-specific. **a**, *gfp::h2b, rde-3(-); gfp::h2b* and WT (no *gfp::h2b*) animals were fed *E.coli* expressing either empty vector control or *gfp* dsRNA. **b,** WT and *rde-3(-)* animals were fed *E.coli* expressing empty vector control and either *oma-1* or *dpy-11* dsRNA. *gfp* (**a**), *dpy-11* and *oma-1* (**b**) pUG RNAs were detected using the assay outlined in Fig. 1a.

**Extended Data Fig. 2.**
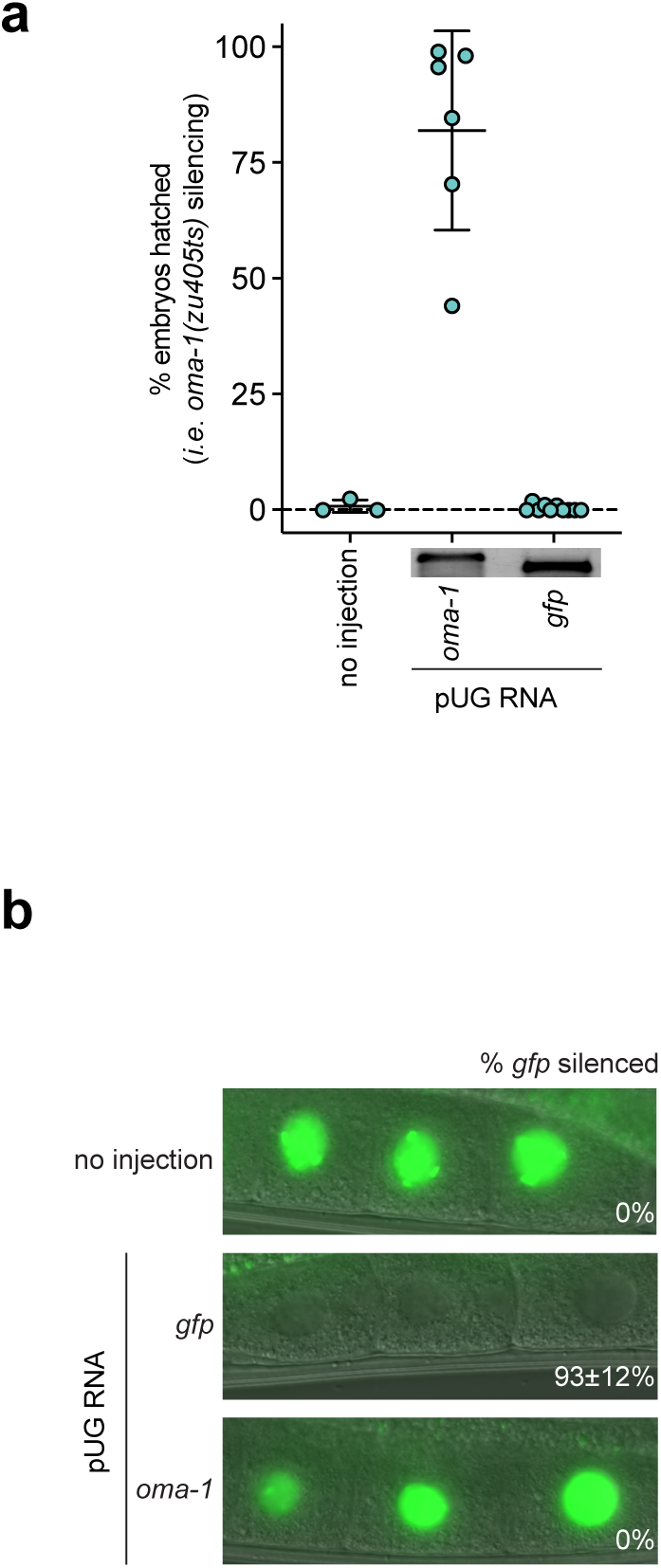
pUG RNA-directed gene silencing is specific. **a**, *rde-1(ne219); oma-1(zu405ts)* animals were injected with either an *oma-1* or *gfp* pUG RNA. Inset shows injected RNAs run on a 2% agarose gel to assess RNA integrity. **b,** *rde-1(ne219); gfp::h2b* animals were injected with either an *oma-1* or *gfp* pUG RNA. % embryonic arrest **(a)** and % gfp silencing **(b)** were scored. All pUG tails were 36nt in length. n = 6-10 injected animals.

**Extended Data Fig. 3.**
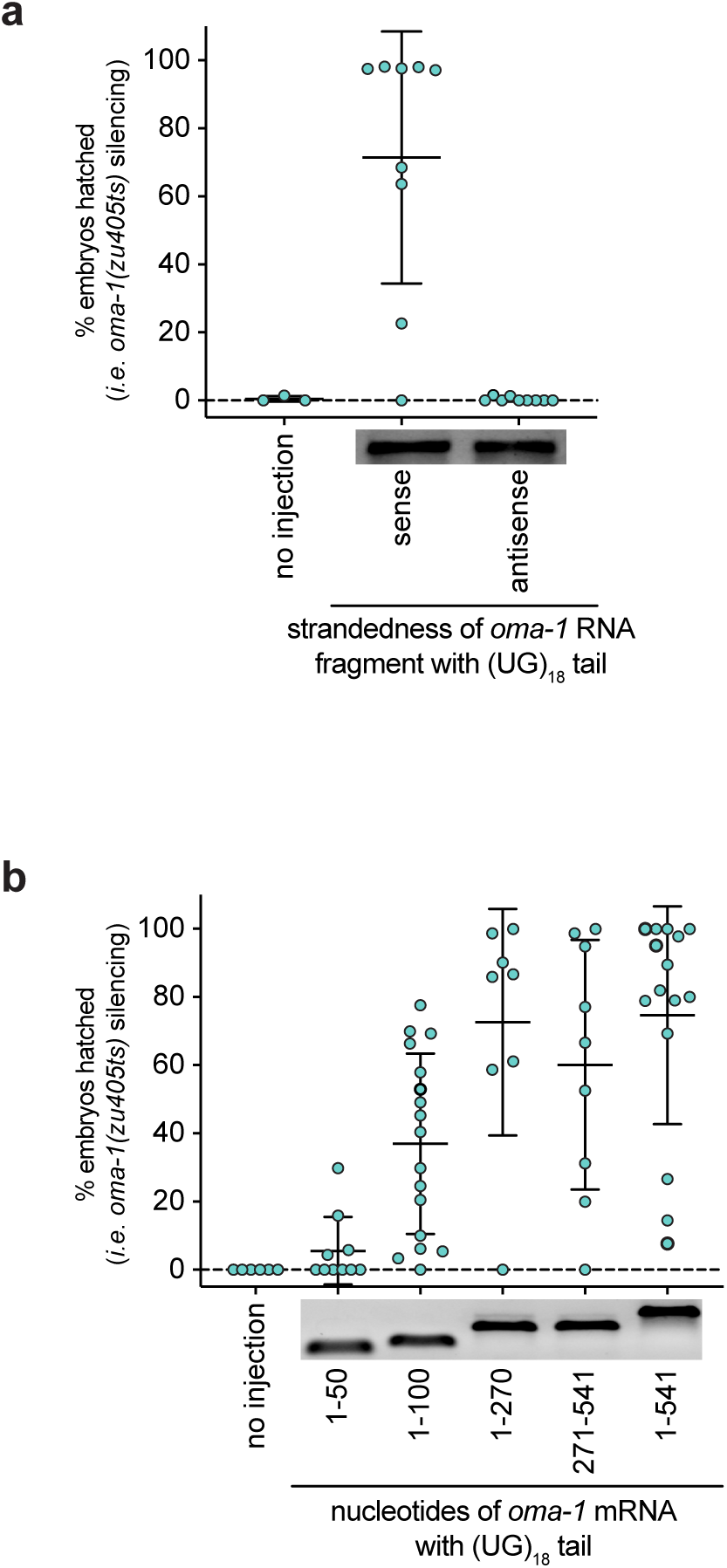
pUG tails must be appended to sense RNAs of >50 nts for functionality. *rde-1(ne219); oma-1(zu405ts)* animals were injected with: **a,** an *oma-1* pUG RNA consisting of the sense or antisense strand of the same 541nt long *oma-1* mRNA fragment (beginning at the *atg)* and a 36nt 3’ pUG tail. **b,** *oma-1* pUG RNAs consisting of *oma-1* mRNA fragments of varying lengths (with position 1 starting at the *aug* of the *oma-1* mRNA sequence) all appended to a 36nt pUG tail. For **a** and **b**, % embryonic arrest was scored at the non-permissive temperature for *oma-1(zu405ts)*. n=8-17 injected animals.

**Extended Data Fig. 4.**
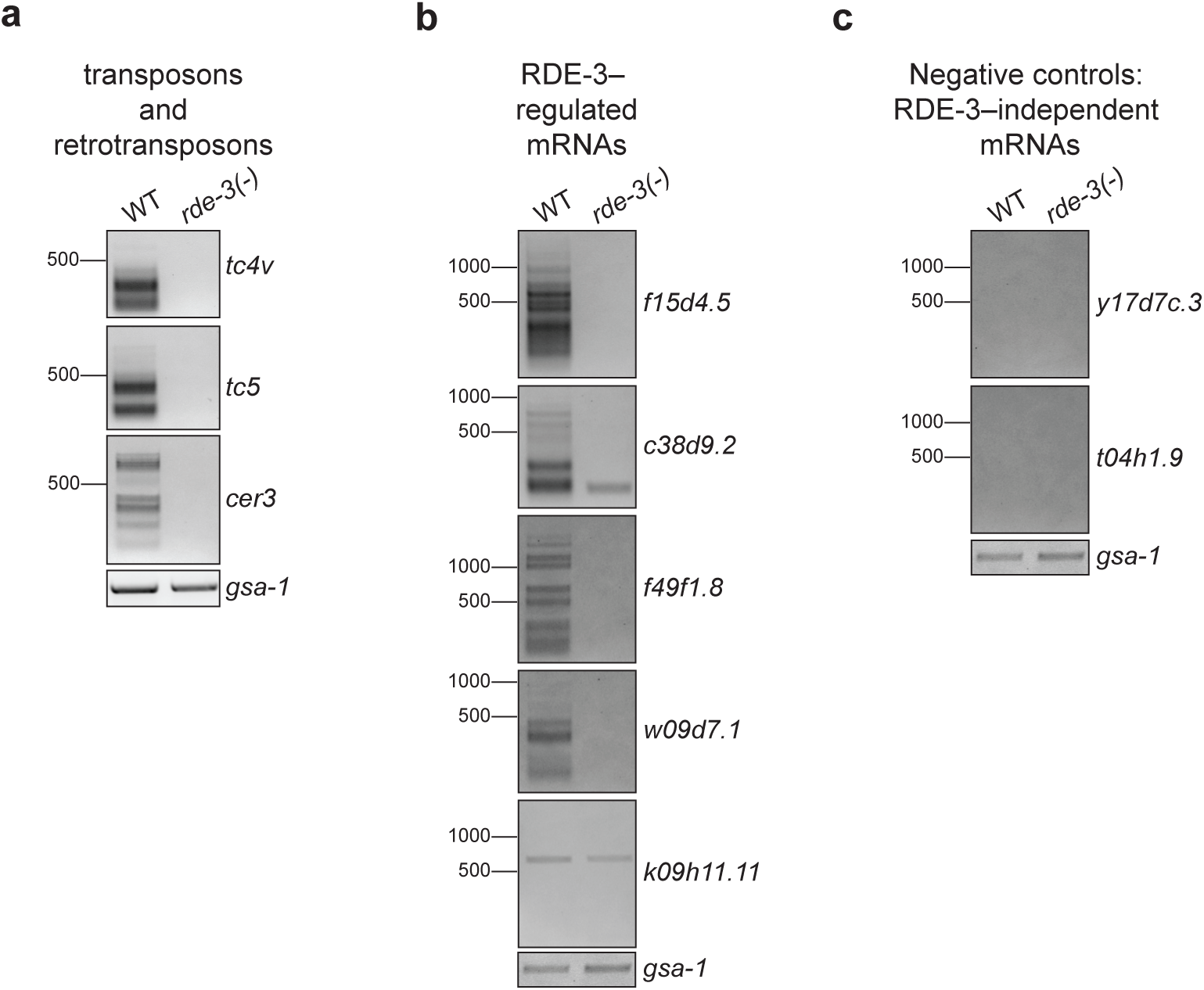
Endogenous targets of pUGylation in *C. elegans*. Total RNA was extracted from WT or *rde-3(-)* animals. The assay outlined in Fig. 1a was used to detect pUG RNAs for **a-b,** RNAs that were significantly upregulated in *rde-3(-)* animals; and **c,** two randomly selected RNAs whose expression does not change in *rde-3(-)* mutants. Data is representative of 3 biological replicates. The same RT samples were used for panels **b** and **c** and, therefore, the *gsa-1* loading control is the same for both panels.

**Extended Data Fig. 5.**
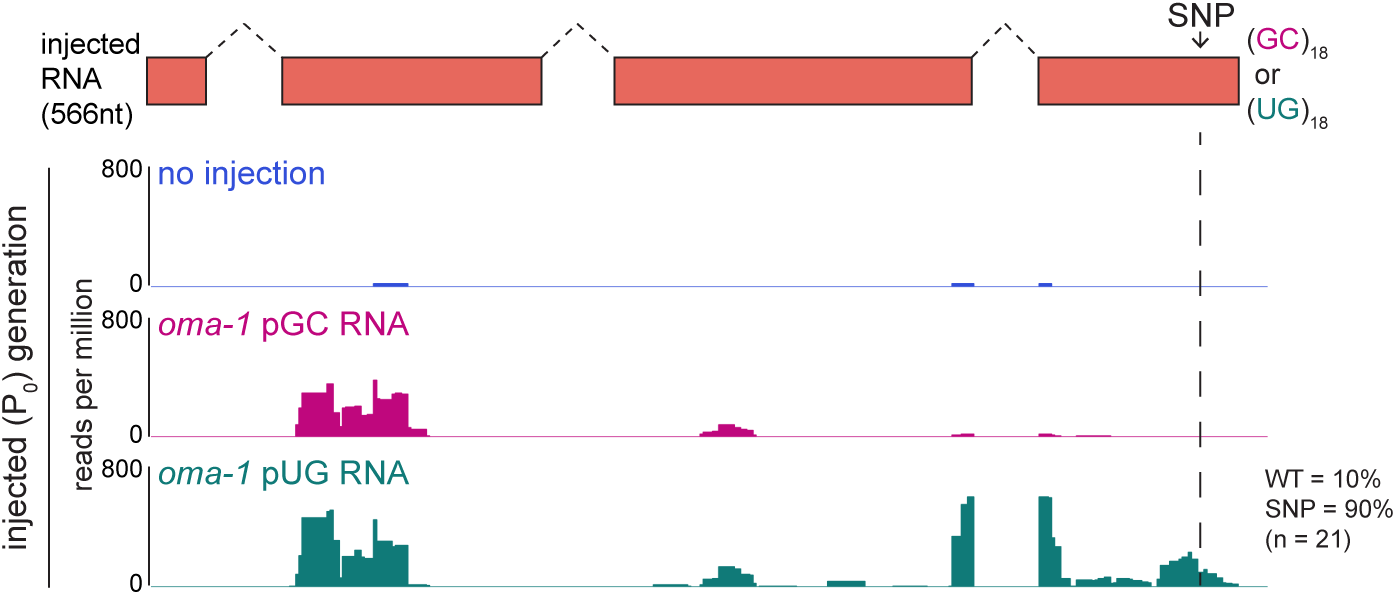
pUG RNAs are templates for RdRPs. A biological replicate of the experiment shown in Fig. 4c was performed. *oma-1(SNP)* pUG or pGC RNAs were injected into *rde-1(ne219); oma-1(zu405ts)* germlines. SNP is indicated with the dotted line. Total RNA was isolated 4-6 hours after injection and small RNAs (20-30nts) were sequenced. 22G siRNAs mapping antisense to *oma-1* are shown. *oma-1* pUG (but not pGC) RNA triggered 22G siRNA production near the site of the pUG tail (pUG-specific 22G siRNAs). For unknown reasons, both pUG and pGC RNA injections triggered small RNA production ≅400bp 5’ of either tail.

**Extended Data Fig. 6.**
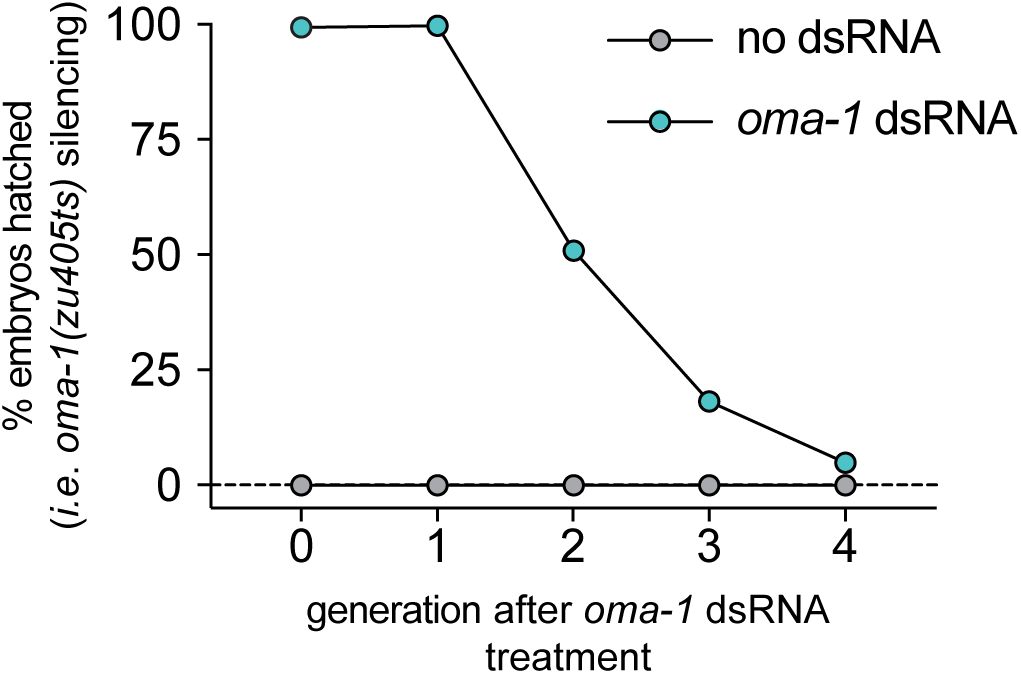
*oma-1* dsRNA triggers heritable silencing. *oma-1(zu405ts)* animals were fed *oma-1* dsRNA and % embryos hatched was scored for 5 generations. Data represents one biological replicate.

**Extended Data Fig. 7.**
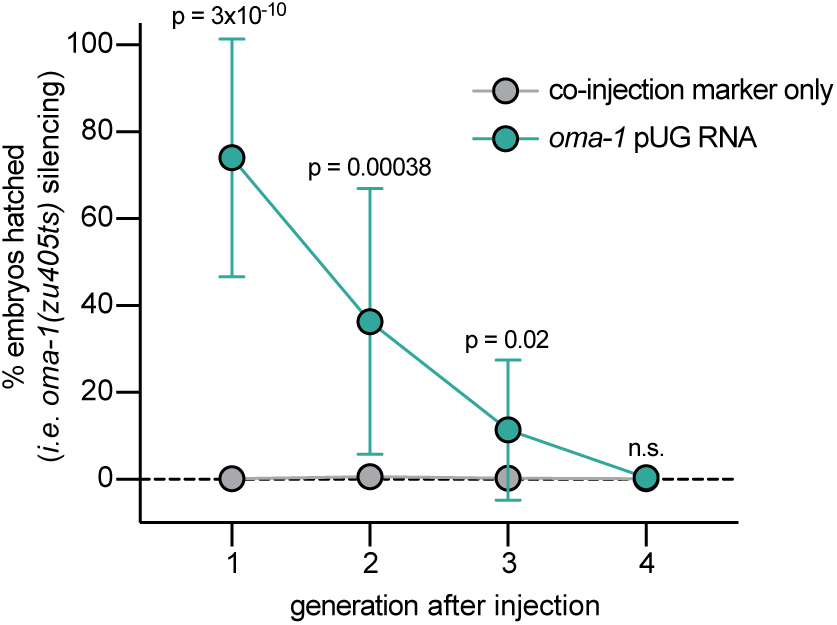
pUG RNA injection triggers heritable silencing. *rde-1(ne219); oma-1(zu405ts)* animals were injected with co-injection marker +/- *oma-1* pUG RNA and % embryonic arrest was scored for four generations in lineages of animals established from injected parents. The data show that *oma-1(zu405ts)* is silenced for multiple generations after an *oma-1* pUG RNA injection.

**Extended Data Fig. 8.**
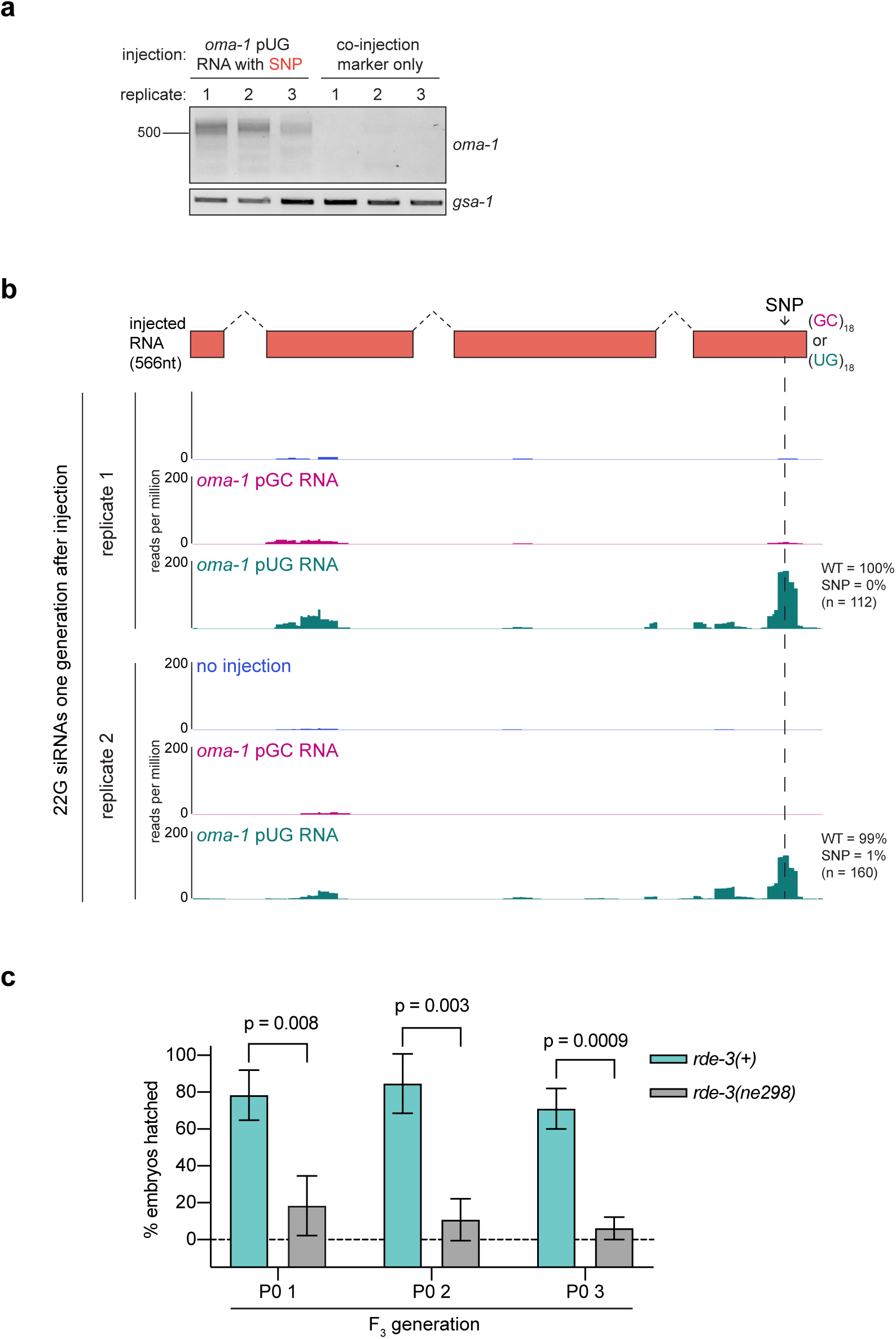
*de novo* pUGylation events in progeny are required for TEI. **a**, *rde-1(ne219); oma-1(zu405ts)* animals were injected with an *oma-1(SNP)* pUG RNA or with co-injection marker only. Co-injection marker-expressing F_1_ progeny were picked and allowed to lay their F_2_ broods. *oma-1* pUG PCR was performed on total RNA from F_2_ progeny. **b,** Two biological replicates of 22G siRNAs sequenced from the progeny of *rde-1(ne219); oma-1(zu405ts)* animals injected with *oma-1(SNP)* pUG or pGC RNAs are shown. Dotted line indicates location of SNP. 22G siRNA reads were normalized to total number of reads. In Fig. 4c and **Fig. S5**, 22G siRNAs were sequenced 4-6 hours after injection and 100% were found to encode the complement of the engineered *oma-1* SNP. Shown here, <1% of 22G siRNAs from progeny of injected animals encoded the SNP complement (insets). Note: siRNAs mapping near the pUG tail were observed only after *oma-1(SNP)* pUG RNA injection (pUG-specific siRNAs). For unknown reasons, both *oma-1(SNP)* pUG and pGC RNAs triggered small RNA production 5’ of the pUG-specific siRNAs. It is possible that these siRNAs were triggered by the piRNA system. Further work will be needed to ascertain the etiology of these RNAs. **c,** *oma-1(zu405ts)* and *rde-3(ne298); oma-1(zu405ts)* animals were fed and mated on *oma-1* dsRNA. F_2_ progeny from this cross were genotyped for *rde-3(ne298)* and F_3_ progeny were phenotyped for % embryonic arrest. 3 biological replicates (P_0_ 1-3) were performed at the non-permissive temperature for *oma-1(zu405ts)*. Error bars are +/- s.d.

**Extended Data Fig. 9.**
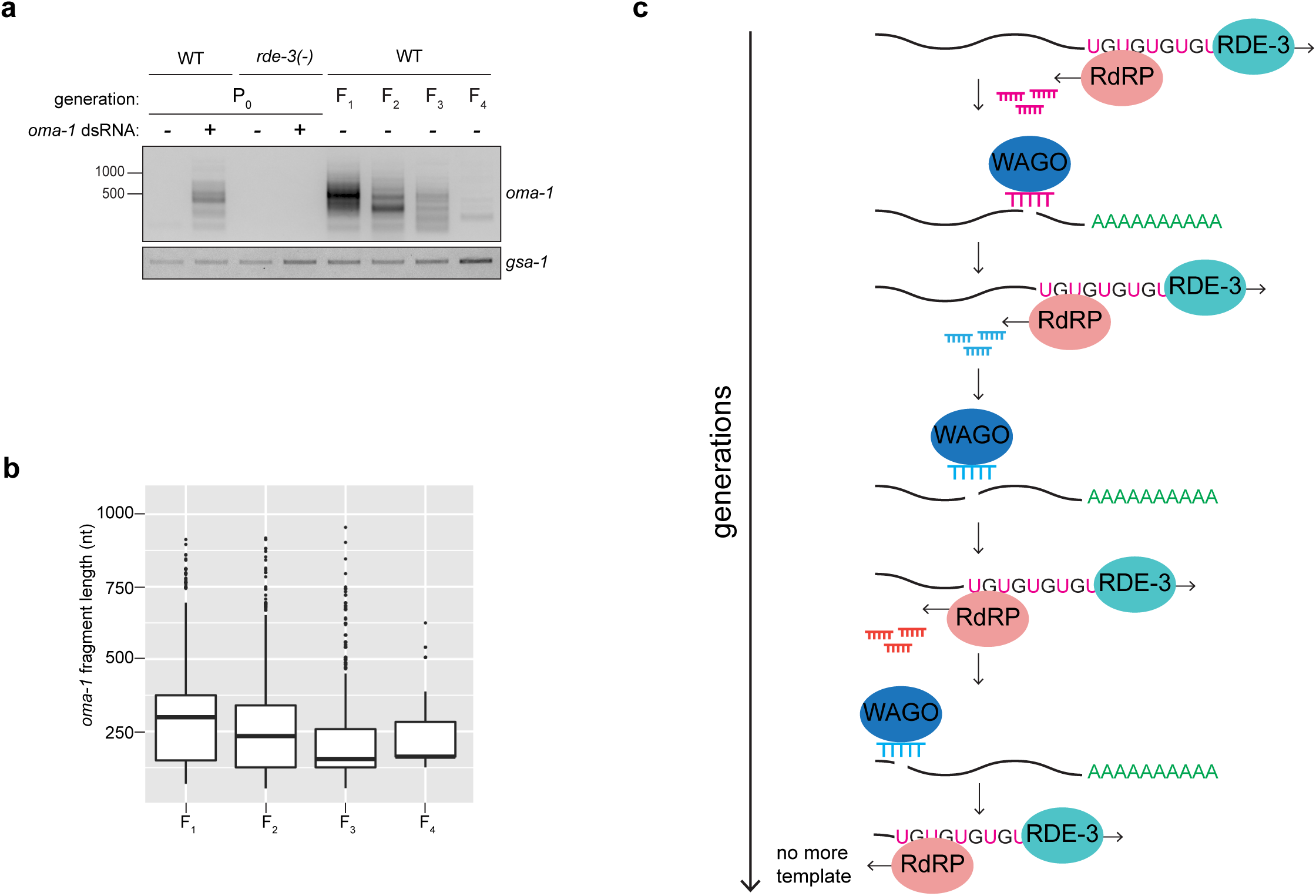
pUG RNA shortening may act as a brake on TEI. **a**, The gel shown is the same as in Fig. 5a, except that *oma-1* pUG RNAs from the P_0_ generation are included. **b,** *oma-1* pUG RNA reads from MiSeq were mapped to *oma-1* and the length of the *oma-1* mRNA portion of each pUG RNA was determined (y-axis). Shown is a Box and Whisker plot representing the interquartile range (IQR, box) and median (line in the box) of lengths at the indicated generations after dsRNA-treatment. The y-axis starts at the *aug* of the *oma-1* mRNA. The whiskers extend to values below and above 1.5*IQR from the first and third quartiles, respectively. Data beyond the end of the whiskers are outliers and plotted as points. The data support the gel in **a**, showing that pUG RNAs get shorter in each generation during RNAi-directed TEI. **c,** A “ratchet” model to explain pUG RNA shortening. pUG RNA shortening may be due to the 3’→5’ directionality of RdRPs, which causes each turn of the pUG/siRNA cycle (see model in Fig. 5g) to trigger cleavage and pUGylation of target mRNAs at sites more 5’ than in the previous cycle until, eventually, pUG RNAs are too short to act as RdRP templates, thereby ending the cycle. Additional support for the ratchet model comes from Fig. 3g and 5c, which show that RNAi-triggered pUG RNAs are longer in *mut-16* and MAGO12 mutant animals than in wild-type animals. Our data indicates that loss of MUT-16 or the WAGOs blocks pUG/siRNA cycling, suggesting that MUT-16 and the WAGOs act during the pUG/siRNA cycling phase of pUG RNA-mediated gene silencing (see model in Fig. 5g). In the absence of cycling, pUG shortening does not occur and pUG RNAs are longer in *mut-16* and MAGO12 animals. Finally, a number of recent studies report transgenerational inheritance of acquired traits in *C. elegans*, which last 3-4 generations^32–37^. *oma-1* RNAi-directed pUG RNAs also perdure for 3-4 generations (Fig. 5a). These shared generational timescales of inheritance suggest that the inheritance of acquired traits in *C. elegans* may be mediated by pUG RNAs whose generational “half-life” is limited to 3-4 generations due to the built-in brake on TEI provided by pUG RNA shortening.

## Supplementary Tables

**Table S1. pUG RNA sequencing data.** This table contains *oma-1* pUG RNA reads from miSeq, our calculations of the accuracy of pUG tails and Tc1 pUG RNAs sequenced using Sanger sequencing.

**Table S2. Genes upregulated in *rde-3* mutants.** List of upregulated RNAs in *rde-3(-)* mutants (adjusted p value <0.05 and log2fold change >1.5).

**Table S3. Small RNA reads mapping to *oma-1.*** *oma-1* small RNAs sequenced after *oma-1(SNP)* pUG and pGC RNA injections (with a no injection control) in either the injected generation or from the progeny of injected animals.

**Table S4. Oligos, *C. elegans* strains and pUG RNAs used in this study.**

## Methods

### Genetics

*C. elegans* culture and genetics were performed as described previously^1^. Unless otherwise noted, all *C. elegans* strains (see Table S4) were maintained at 20°C on NGM growth media and fed OP50 bacteria.

### RNAi

Embryos were obtained via hypochlorite treatment of gravid adult hermaphrodites (egg prep) and dropped onto plates seeded with HT115 bacteria expressing dsRNA against a gene of interest. After 3-4 days, gravid adults were washed off plates using M9 + Triton X-100 buffer, collected in Trizol, flash frozen in liquid nitrogen and stored at -80°C until total RNA extraction. The *dpy-11* and one of the *oma-1* RNAi clones came from the *C. elegans* RNAi collection (Ahringer lab). The second *oma-1* RNAi clone was a custom clone made to target exon 6 of *oma-1*. The *gfp* RNAi clone was obtained from the Fire lab. For transgenerational RNAi inheritance experiments, embryos were seeded onto plates with either empty vector control or dsRNA of interest. Some gravid adults were collected for the P_0_ generation sample and the remaining were egg prepped onto plates without dsRNA every generation, for the indicated number of generations.

### pUG PCRs and qPCRs

Total RNA was extracted using TRIzol Reagent (Life Technologies, 15596018). 5ug of total RNA and 1pmol of reverse transcription oligo was used to generate first-strand cDNA using the Superscript III First-Strand Synthesis System (Invitrogen, 18080051). 1ul of cDNA was used for the first PCR (20ul volume) performed with Taq DNA polymerase (New England BioLabs, M0273) and primers listed in Table S4. First PCR reactions were diluted 1:100 and then 1ul was used for a second PCR (50ul volume) using primers listed in Table S4. PCR reactions were then run on agarose gels. For Sanger sequencing, lanes of interest were cut out from agarose gels and gel extracted using a QIAquick Gel Extraction Kit (Qiagen, 28706). 3ul of gel extracted PCR product was used for TA cloning with the pGEM®-T Easy Vector System (Promega, A1360) according to manufacturer’s instructions. Ligation reactions were incubated overnight at 4°C. Transformations were performed with 5-alpha Competent *E. coli* cells (NEB, C2987H) and plated on LB/ampicillin/IPTG/X-Gal plates. White colonies were selected, inoculated and miniprepped using QIAprep Spin Miniprep Kit (Qiagen, 27106). Plasmids were sequenced at the Dana-Farber/Harvard Cancer Center DNA Resource Core using a universal SP6 primer (5’-CATACGATTTAGGTGACACTATAG-3’). qPCRs were performed using 2ul of 1:100 diluted first PCRs as a template with qPCR primers (Table S4) and iTaq Universal SYBR Green Supermix (Bio-Rad) according to manufacturer’s instructions.

### MiSeq

*oma-1* pUG PCRs were sequenced on an Illumina MiSeq from animals fed HT115 bacteria expressing empty vector control plasmid, *oma-1* dsRNA from the Ahringer RNAi library or our custom *oma-1* dsRNA (P_0_-F_4_ generations for this experiment). A first round of PCR was performed with the same primers as described above. Primers were modified for the second PCR to contain Illumina p5 and p7 sequences, read 1 and 2 sequencing primers, a unique index (reverse primer only) for multiplexing and unique molecular identifiers (NNN) (Table S4). PCR reactions were then pooled, run on an agarose gel and gel purified as described above. Sequencing was performed on an Illumina MiSeq to obtain paired-end reads (67bp for Read 1, 248bp for Read 2).

### MiSeq sequencing analysis

First, unique molecular identifiers (UMIs) were removed from each read pair and appended to the end of the read name using UMI-tools^2^. Then, cutadapt v2.5 was used for the following: 1) low-quality bases (quality score < 20) were trimmed from the 3’ ends of reads; 2) read pairs containing the inline portion of the 5’ adapter (AACAACGAGAAGATCGATGA) in Read 1 were selected for and then trimmed; 3) Read pairs containing the inline portion of the 3’ adapter (GGCGTCGCCATATTCTACTTACACACACACACACACAC) in Read 2 were selected for and trimmed; and 4) If the 5’ adapter was present in any Read 2 sequences, the adapter was trimmed from those sequences^3^. After adapter trimming, Read 2 sequences were screened for additional pACs at the 5’ end: reads that did not contain additional pACs (and therefore did not have a pUG tail longer than the adapter) were discarded; reads that did contain additional pACs were retained, and the pACs were trimmed using cutadapt v2.5 (pAC and pCA sequences were provided as non-internal 5’ adapters)^3^. After pAC trimming, Read 2 sequences shorter than 5 nucleotides were discarded. The remaining Read 2 sequences were aligned to the *C. elegans* genome (WormBase release WS260) using STAR v2.7.0f ^4^. SAM and BED files of unique alignments were generated using SAMtools v1.9 and BEDtools v2.27.1 and then imported into R for subsequent analyses^5–7^. Alignments were deduplicated based on the combination of the UMI and end coordinate. Alignments that mapped to the “+” strand and/or to coordinates outside of the *oma-1* gene were discarded.

To systematically define the “*oma-1*” and “pUG” portions of each read, the pre-pAC-trimmed version of the read was reverse-complemented and then split as follows. By default, the aligned portion of the read was designated as “*oma-1*”, and any sequence downstream of the aligned portion was designated as the “pUG.” Then, the “*oma-1*” portion was matched to an *oma-1* reference sequence (spliced + UTRs) using Biostrings v2.50.2^8^. If the first 1-6 nucleotides that occurred 3’ of the match were the same in the *oma-1* reference as they were in the read prior to pAC trimming (and therefore had the potential to be templated), then those nucleotides were reassigned to the “*oma-1*” portion of the read. End coordinates of the alignments were adjusted accordingly. A small portion of reads (<15%) were misannotated with the above approach, largely due to soft-clipping at the 3’ end during alignment. To systematically filter out such reads, reads for which the annotated “pUG” started with a base other than “U” or “G” and/or contained 2 or more bases other than “U” or “G” within the “pUG” sequence were discarded. The abundance of each pUGylation site (Figure 1e) was plotted in R using Sushi v1.20.0^9^. To generate the pUG site logos shown in Figure 1f, a list of unique pUG sites was sorted by the last nucleotide of the “*oma-1*” portion and then plotted in R using ggseqlogo v0.1^10^.

### pUG RNA injections

*<u>gfp</u>* <u>and *oma-1* pUG RNAs.</u> pUG RNAs were synthesized *in vitro* using MEGAscript T7 Transcription Kit (Invitrogen, AM1334). DNA templates for *in vitro* transcription reactions were gel purified PCR products amplified using primers listed in Table S4. 150ng of gel purified PCR products was used as a template. Reactions were incubated overnight at 37°C. *in vitro* transcribed RNA was purified using TRIzol Reagent (Life Technologies, 15596018) and stored at -80°C. Injection mix consisted of 0.5pmol/ul *in vitro* transcribed RNA and 2.5ng/ul co-injection marker (*pmyo-2::mCherry::unc-54 3’UTR*) plasmid pCFJ90 (Addgene, plasmid #19327), dissolved in water. Animals expressing co-injection marker show *mcherry* expression in the pharynx. Adult hermaphrodites ((either *gfp::h2b; rde-1(ne219)* for *gfp* pUG RNA injections or *oma-1(zu405); rde-1(ne219*) for *oma-1* pUG RNA injections) were injected in the germline and allowed to recover at 15°C for two days before being shifted back to 20°C. Adult progeny of injected animals expressing *mcherry* in the pharynx were picked under an Axis Zoom.V16 fluorescent dissecting microscope using a PlanNeoFluar Z 1x/0.25 FWD 56mm objective and scored *gfp* or *oma-1* expression. *gfp* expression was scored using the Plan-Apochromat 20 × /0.8 M27 objective on an Axio Observer.Z1 fluorescent microscope (Zeiss). Images were taken with the Plan-Apochromat 63 × /1.4 Oil DIC M27 objective. *oma-1(zu405ts*) is a gain-of-function temperature-sensitive allele of *oma-1*. *oma-1(zu405ts)* animals lay arrested embryos at 20°C^11^, unless *oma-1(zu405ts)* is silenced. To measure *oma-1(zu405ts)* silencing, five progeny from each injected animal were transferred to a new plate. Animals were removed after laying 50-100 eggs, and *oma-1(zu405)* silencing was measured as percentage of eggs hatched. <u>Tc1 pUG RNA</u>. T7 *in vitro* transcription was performed as described above to synthesize a Tc1 pUG RNA consisting of a 36nt pUG tail appended to a 338nt long fragment of Tc1 RNA (see Table S4 for primers used). This Tc1 pUG RNA was injected (as above) into the germlines of *rde-3(ne3370); unc-22(st136)* animals. *unc-22(st136)* animals have a Tc1 DNA transposon insertion in the *unc-22* gene, resulting is paralysis. Co-injection marker-expressing progeny of injected animals were picked at the L4 stage and pooled (25 animals per pool) onto 10cM NGM plates and allowed to lay a brood. The number of mobile adult progeny in each pool was counted 6-7 days later.

### RNA FISH + Immunofluorescence

Approximately 30 animals were dissected in 15 μl of 1X egg buffer (25 mM HEPES (pH 7.3), 118 mM NaCl_2_, 48 mM KCl, 2 mM CaCl_2_, 2 mM MgCl_2_) to isolate gonads. A coverslip was placed on top of dissected tissue, excess buffer was soaked up using a Kimwipe and slides were placed onto a metal block pre-chilled on dry ice for 10 min. Coverslips were popped off and slides were submerged in methanol at −20°C for 10 min. Slides were then washed twice, 5 min per wash in 1X PBS + 0.1% Tween-20 (PBSTW). Samples were then fixed with 4% paraformaldehyde solution in 1X PBS for 20 minutes, followed by two 5 min washes in PBSTW. Samples were then incubated at 37°C for 6 hours in a humid chamber with a 1:50 dilution of fluorescent RNA FISH probe in hybridization buffer (10% formamide, 2X SSC, 10% dextran sulfate (w/v)). The RNA FISH probe to detect pUG RNAs (/5Alex647N/CACACACACACACACACACA) was ordered from Integrated DNA Technologies (IDT) and stored at a stock concentration of 100uM at -20°C. The RNA FISH probe to detect *ama-1* mRNA was ordered from Stellaris (SMF-6011-1). After 6 hours, slides were washed twice, 10 min per wash, in FISH Wash Buffer (2X SSC, 10% formamide, 0.1% Tween-20). Samples were then washed for 5 min in 2X SSC. Slides were sealed using 15ul of Slowfade Gold with DAPI. For experiments in which RNA FISH and immunofluorescence were combined, RNA FISH was first performed as above. After the final 2X SSC wash, slides were washed once with PBST for 5 min, samples were incubated overnight at room temperature in a humid chamber with a 1:1000 dilution of GFP antibody (Abcam, ab290) in PBSTW. Slides were then washed three times, 10 min per wash, in PBSTW and incubated in a 1:100 dilution of Alexa Fluor 488 goat anti-rabbit in PBSTW for 2 hours at room temperature in a humid chamber. Slides were next washed three times, 10 min per wash, in PBSTW and then sealed with 15ul of Slowfade Gold with DAPI. All imaging was performed on an Axio Observer.Z1 fluorescent microscope (Zeiss) using the Plan-Apochromat 63 × /1.4 Oil DIC M27 objective. All image processing was done on Fiji^12^.

### RNA-seq

Total RNA was extracted using TRIzol Reagent (Life Technologies, 15596018). RNA quality (RIN) and quantity were assessed on the TapeStation 2200 (Agilent). Two rounds of mRNA purification were performed on 1ug total RNA using the Dynabeads mRNA DIRECT Kit (Invitrogen, 61011). First-strand cDNA was generated using the Superscript III First-Strand Synthesis System (Invitrogen, 18080051), followed by second-strand synthesis using DNA polymerase I (Invitrogen, 11917010). cDNA libraries were prepared using the Nextera XT DNA Library Preparation Kit (Illumina, FC-131-1024). Libraries were sequenced on the Illumina NextSeq500 platform (Biopolymers Facility, HMS) and 75 bp paired-end reads were obtained.

### RNA-seq analysis

Reads were trimmed to remove sequencing adapters and low-quality bases using Trim Galore version 0.4.4_dev (https://www.bioinformatics.babraham.ac.uk/projects/trim_galore/). Trimmed reads were then aligned to the *C. elegans* genome (UCSC ce11/WBcel235) using STAR version 2.7.0a^13^. Differential expression analysis of genes and repeat elements was performed using the TEtranscripts package in TEToolkit version 2.0.3^14^. Gene annotations were obtained from Ensembl (WormBase release WS260)^15^. Repeat annotations were obtained from UCSC by downloading the RepeatMasker (rmsk) table in the Table Browser program. The table was reformatted to a GTF file using the Perl script makeTEgtf.pl (http://labshare.cshl.edu/shares/mhammelllab/www-data/TEToolkit/TE_GTF/). Features with an adjusted p value of < 0.05 and a log2 fold change > 1.5 were reported.

### CRISPR

The CRISPR strategy described previously^16^ was used to revert the missense mutation in *rde-3(ne298)* animals to wild-type and to tag the N-terminus of *rrf-1* with *ha::tagRFP*. SapTrap cloning^17,18^ and the selection-based CRISPR strategy described previously^19^ was used to introduce a *gfp* tag at the C-terminus of *c38d9.2*, to tag *rde-3* at the N-terminus with *gfp::degron* and to introduce *3xflag::rde-3* (with 2kb upstream of the ATG and 2kb downstream of the stop codon) at the LGII MosSCI site *ttTi5605*^20^ into *rde-3(ne3370)* animals. All guide RNAs were designed using the guide RNA selection tool CRISPOR^21^.

### Small RNA sequencing

*rde-1(ne219); oma-1(zu405)* animals were injected with an *oma-1* pUG or pGC RNA in which the *oma-1* sequence (the first 566nt of *oma-1* mRNA) was modified to contain a SNP in exon 4 (ATTCATCCCG A>T TCATGGACCA). Injection mix was prepared as described above. For P_0_ analysis, ∼100 *rde-1(ne219)* animals were injected per experiment. After recovering for 1-4 hours at room temperature, injected animals were collected for total RNA extraction. For F_1_ analysis, ∼20 *rde-1(ne219)* animals were injected per experiment. Injected animals recovered at 15°C for two days and were returned to room temperature. ∼500 adult co-injection marker-expressing progeny of injected animals were collected for total RNA extraction. Small RNAs were size-selected, cloned and sequenced as described previously^22^. Note: the same SNP-containing *oma-1* pUG RNA was used for Figure 5d, in which co-injection marker-expressing progeny of injected animals were picked and allowed to lay a brood that was collected for *oma-1* pUG PCR analysis as described above.

### Small RNA sequencing analysis

A custom Python script was used to select reads starting with the last 4 nucleotides of the 5’ adaptor (either AGCG or CGTC). Cutadapt 1.14^3^ was then used to trim the 3’ adaptor (CTGTAGGCACCATCAATAGATCGGAAGAGCAC) and the in-line portion of the 5’ adaptor (AGCG and CGTC) (both with a minimum phred score = 20), allowing only sequences >= 16nt after trimming to pass (cutadapt -q 20 -m 16 -u 4 -a CTGTAGGCACCATCAATAGATCGGAAGAGCAC --discard-untrimmed). The quality of the trimming was assessed with FastQC 0.11.5^23^. For downstream analysis, custom Python scripts were used to select reads that were 22nt in length and began with a G (22G siRNA reads). Tophat 2.1.1^24^ was then used to map 22G siRNA reads to the *C. elegans* genome (WBcel235). Gene annotations were obtained from Ensembl^15^ (WormBase release WS269) and custom shell scripts were used to select protein-coding genes only. One mismatch was allowed to identify 22G siRNAs with SNPs. Using Samtools v0.1.19, only uniquely mapping sequences were retained. 22G RNA pileup figures were generated as follows: first, bam files generated from Tophat v2.1.1^24^ were normalized by DeepTools v3.0.2^25^ based on counts per million and only antisense reads were kept for further analysis (bamCoverage -bs 2 --normalizeUsing CPM -samFlagExclide 16). Then, the normalized antisense 22G small RNA sequences (bedGraph files) were visualized using Sushi 1.20.0^9^ in R. The number of reads mapping antisense to each gene was calculated by featureCounts 1.6.0^26^ (featureCounts -s 2 -a *.gtf -t exon -g gene_name). All custom scripts used in this section are available at: https://github.com/Yuhan-Fei/pUG-analysis.

### pUG RNA chromatography

Adult animals (∼1-2 full 10cm plates per experiment) were frozen in liquid nitrogen as small droplets and ground into powder with a mortar and pestle. Powder was dissolved in lysis buffer (5mM HEPES-NaOH(pH7.5), 50mM NaCl, 5mM MgCl_2_, 0.5mM EDTA (pH8.0), 5% glycerol, 0.25% Triton X-100, 0.5mM DTT, 1mM PMSF, 1 tablet of cOmplete protease inhibitor (Roche, 11697498001)) and rotated for 30 minutes at 4°C. The resulting lysate was centrifuged at top speed for 10 min at 4°C. Supernatant was distributed evenly among experiments, and RNaseOUT recombinant ribonuclease inhibitor (Invitrogen, 10777019) was added to lysate (1ul per 100ul lysate). For each experiment, 160pmol of biotinylated RNA was conjugated to 400ug Dynabeads MyOne Streptavidin beads (Invitrogen, 65001) as per manufacturer’s instructions. Beads were added to lysates and rotated at room temperature for 1 hour. Beads were separated from supernatant on a magnetic rack, and the supernatant was collected and saved (“sup” fraction). Beads were washed 3 times with lysis buffer and rotated for 5 min at 4°C in lysis buffer. Supernatant was removed, beads were dissolved in 1x Laemmli sample buffer (Biorad, 1610737) with 5% 2-mercaptoethanol and heated for 5 min at 95°C and chilled on ice (“pull-down” fraction).

### Gel electrophoresis and Western blot

“Pull-down” and “sup” fractions were loaded into 4–15% Mini-PROTEAN TGX Precast protein gels (Biorad, 4561086) and run in Tris-glycine running buffer (25mM Tris, 192mM glycine, 0.1% SDS). Proteins were then transferred to nitrocellulose membrane (BioRad) at 100V for 1 hour in electrotransfer buffer (50mM Tris, 40mM glycine, 9% methanol, 0.2% SDS). Blotted membranes were blocked with 5% milk in PBST (phosphate-buffered saline, 1.0% Tween-20) for 1 hour at room temperature and probed with primary antibody (1:1000 HA-Tag Rabbit mAb, Cell Signaling, #3724, in 5% milk) overnight at 4°C. After washing with PBST 3 times, membrane was probed with secondary antibody (1:10,000 IRDye 800CW Goat anti-Rabbit IgG, LI-COR, 926-32211, in 5% milk) for 1 hour at room temperature. Membrane was washed with PBST 3 times before imaging using Odyssey Fc Dual-Mode Imaging System (LI-COR).

### Heterozygous Experiment

Embryos were obtained via hypochlorite treatment of wild-type gravid adult hermaphrodites and dropped onto plates seeded with HT115 bacteria expressing dsRNA targeting *oma-1.* L_4_ hermaphrodites were then transferred, along with *rde-3(ne298)* males, onto “mating plates” seeded with 25ul of *oma-1* dsRNA-expressing bacteria. Once hermaphrodites were adults, they were singled onto NGM plates seeded with OP50 and allowed to lay a brood. 12-15 F_1_s were singled from 3 independently mated hermaphrodites and genotyped to ensure that they were heterozygous for *rde-3(ne298).* To obtain F_3_ animals, 12-15 F_2_s per F_1_ were singled to 15°C (so as to avoid embryonic arrest due to temperature) and allowed to lay a brood. F2s were then single worm genotyped to identify *rde-3(+)* and *rde(ne298)* homozygous animals. Then, % embryonic arrest was calculated by pooling 5 L_4_ stage F_3_ animals per F_2_ at 20°C until they had laid a brood of 50-200 progeny and counting the # of embryos that were laid vs. hatched on the following day. *rde-3(+)* and *rde(ne298)* homozygous F_3_ broods were pooled for all plates that were derived from the same P_0_ and pUG PCR was performed as described above.

## Acknowledgements

We thank members of the Kennedy, Butcher and Wickens labs for helpful discussions; the Biopolymers Facility at HMS for Illumina sequencing; and the Dana-Farber/Harvard Cancer Center DNA Resource Core for Sanger sequencing. Some strains were provided by the *Caenorhabditis* Genetics Center (CGC), which is funded by the NIH Office of Research Infrastructure Programs (P40 OD010440). Some strains were provided by the Mitani laboratory through the National BioResource Project (Tokyo, Japan), which is part of the International *C. elegans* Gene Knockout Consortium.

## Author contributions

A.S. contributed to Figs. 1a-f, 3a-g, 5a, 5c-g; Extended Data Figs. 1a-c; 4a-c; 6; 8a, 8c; 9a-c; and Supplementary Tables S1, S2, S4. J.Y. contributed to Figs. 2a-c, 3h, 4a-c, 5b; Extended Data Figs. 2a-b, 3a-b; 5, 7, 8b; Supplementary TableS S3, S4. D.J.P., J.G. and J.G.S. contributed to Extended Data Fig. 4 and Supplementary Table S2. A.E.D. contributed to Figs. 1e-f; Extended Data Fig. 9b; and Supplementary Table S1. Y.F. contributed to Fig. 4c; Extended Data Figs. 5, 8b; and Supplementary Table S3. A.S., M.W. and S.K. conceived the project. S.K. supervised the project. A.S. and S.K. wrote the manuscript.

## Competing interests

The authors declare no competing interests.

